# Co-occurrence networks can preserve emergent properties of ecological communities

**DOI:** 10.64898/2026.04.15.718781

**Authors:** Fiona Callahan, Claire Evensen

**Affiliations:** Center for Computational Biology, University of California, Berkeley, United States; Department of Integrative Biology, University of California, Berkeley, United States

## Abstract

Interaction networks, in which nodes represent species and edges represent direct interactions between species, have a long and impactful history in community ecology. However, co-occurrence networks, where edges represent statistical relationships among species presences or abundances, are often easier to construct from lab and field data. It is clear that co-occurrence edges often do not represent direct interactions, but frameworks for the interpretation of co-occurrence networks have not kept pace with their generation. It is therefore unclear when and how these networks can be used to gain insight into community dynamics. Here, we use a Generalized Lotka-Volterra-based model to explore the contexts in which emergent properties of species interaction networks are identifiable in their resulting co-occurrence networks. We find that, in spite of many differences in direct edges, key features of the true interaction network, such as unipartite modularity, high-degree nodes (hubs), and bipartite modularity and nestedness, can be preserved in co-occurrence networks. In contrast, node degree distributions are not preserved even in the most idealized scenarios. We propose that networks derived from large co-occurrence datasets could therefore be used in future empirical work to test existing hypotheses of how emergent network structures drive ecological community dynamics.

## 1 Introduction

Ecological network analysis has a rich history, driven in particular by early developments in food web ecology [1–4]. Multi-omics data collection is now providing new methods to build ecological networks [5], but the underlying motivation remains consistent with the earliest food web studies: characterize patterns of relationships between species to understand the ecosystem as a whole. Studies of complex systems reveal network architectures common across many disparate social, technological, and biological systems [6–9]. Emergent properties of these networks govern core dynamical behaviors of complex systems, such as synchronization, robustness to perturbation, and formation of chimera states [10–13] and have the potential to provide insight into the corresponding processes in complex ecological systems [14–16].

To build a network representation of an ecological community, meaningful nodes and edges must be assigned. Often, nodes represent species, and edges represent direct species interactions (e.g. trophic interactions, mutualisms, host-parasite interactions, etc.). Biotic interactions are one of the major factors that shape species assemblages, and are a central focus of many ecological network studies [17]. However, direct interactions are frequently difficult or – in the case of long-extinct populations – impossible to observe in the lab and field. In such cases, it is common to use spatial or temporal co-abundance or co-occurrence data to construct a co-occurrence network for the ecological community. In this case, the nodes in the network represent taxa and the edges represent statistical relationships between presence or abundance of the observed taxa, such as pairwise correlations. The availability of co-occurrence datasets at a large spatial, temporal, and taxonomic breath is increasing, especially in the era of environmental DNA (eDNA) metagenomics. These data have the potential to revolutionize the study of ecological communities, both in the past and present [18]. However, the ecological interpretation of edges in co-occurrence networks can be complex; though biotic interactions may influence statistical associations between species, they do not always indicate a direct interaction between them [19–21]. For example, species may be associated through indirect interactions mediated through shared interacting partners, or shared responses to the abiotic environment [19]. Some methods attempt to differentiate between direct and indirect interactions in co-occurrence networks, but the degree to which this is successful remains controversial [22–24].

Although problems with inferring accurate direct interactions from co-occurrence data have been well-documented, little work has considered the imprint that broader topological properties of interaction networks may leave on co-occurrence networks [25, 26]. It is these emergent, community-level properties that are hypothesized to drive important ecological processes [27], including local community stability and resilience [28, 29], susceptibility to cascading species extinctions [30], restoration of plant-pollinator interactions [31], and host-pathogen coevolutionary dynamics [32–34]. Applications of ecological networks often fall on the side of descriptive analyses, rather than hypothesis testing, in part due to difficulties in bridging the gap between theoretical and empirical systems [5]. Many of the theoretical advancements in ecological network dynamics are based on models in which pairwise interactions are known precisely, but conclude with hypotheses centered on emergent properties of the network [35]. It stands to reason that if these emergent properties could be determined from co-occurrence data, some of the gaps between theory and experiment could be bridged without perfectly faithful reconstructions of interaction networks.

In this paper we therefore develop a simulation framework to examine which emergent properties of interaction networks are identifiable from co-occurrence networks. Interaction networks of various architectures are constructed for regional species pools, providing a ‘ground truth’ of direct interactions in the ecological community. We then simulate a random dispersal process of community assembly, and use a Generalized Lotka-Volterra (GLV) system of differential equations to simulate population dynamics in the presence of biotic interactions [36]. The abundances of each species at equilibrium in each subsampled community are used – without knowledge of the underlying interaction network – to construct a co-occurrence network among all species in the regional pool. Co-occurrence network properties are then compared to those in the underlying interaction network. Similar methods have been used previously to investigate detection of pairwise species interactions and keystone species [37].

Here, we investigate which properties are preserved between interaction and co-occurrence networks under idealized conditions: perfect measurement of absolute species abundances, spatially independent samples of populations at equilibrium, and (in most cases) no influence of the abiotic environment. We find that even in this framework, many significant co-occurrence associations do not represent direct interactions. However, we find many contexts in which broader network architectures, including relative ordering of generalist and specialist species, unipartite and bipartite modularity, and bipartite nestedness, are robust to these differences. We therefore suggest reframing co-occurrence data as a tool for understanding overall ecological community function, rather than discovering direct ecological interactions.

## 2 Methods

Each iteration of our simulation consists of the following steps: (1) interaction network generation, (2) population dynamics simulations and co-occurrence network generation, and (3) analysis of relevant network metrics. We specifically identify cases when step (1) is not an independent network construction, but rather a systematic perturbation of an initial interaction network. Variables and parameters are defined with their default values in Table S1.

### 2.1 Simulating co-abundance from interactions

At the beginning of each iteration, a Lotka-Volterra community matrix ***A*** was created for the full community of *S* species:

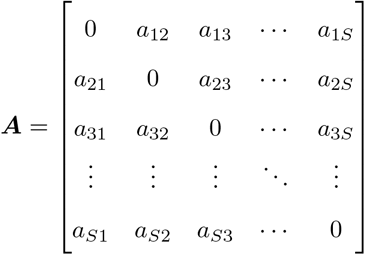

The interaction coefficients *a*_*ij*_, *j* ≠ *i*, specify the effect of species *j* on the population growth of species *i*. Thus, positive values indicate a positive influence on overall growth, and vice versa. This represents a static biotic niche of all of the species, from which sub-networks may form to create realized communities. Interactions were chosen using methods specified below for different network structures.

For each subsampling event, *S*_*m*_ nodes were sampled uniformly at random from the interaction network. This represents a random community assembly process where the species are all equally likely to arrive at a particular sampled location, without priority effects. Abundances were simulated by running a GLV model (for only the subsampled nodes) to equilibrium (see also S1.3).

GLV dynamics of the system were defined as follows:

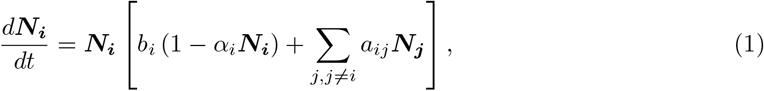

where ***N***_***i***_(*t*) denotes the abundance of species *i, b*_*i*_ is the intrinsic growth rate of species *i*, and *α*_*i*_ is the self-limitation parameter (crowding effects due to intraspecies interactions). The interaction coefficient *a*_*ij*_ is the (*i, j*) entry in the community interaction matrix ***A***.

For an existing interaction, the interaction coefficient was chosen uniformly between 0 and *I*_*max*_, and the sign of the interaction was chosen with probability *q* of being positive. *I*_*max*_ is defined as the product of the absolute value of the intra-species population limiting parameter (*α*_*i*_) and *F*_*max*_, the interaction strength factor. This means, for example, that if *F*_*max*_ = 1/5, the intra-species effect is 5 times stronger than the maximum individual inter-species effect. A vector of intrinsic growth rates ***b*** = [*b*_1_, …, *b*_*n*_] was generated for the full community with each *b*_*i*_ chosen uniformly between 0.9 and 1.

This process was repeated *M* times to simulate *M* spatial samples. For most simulations, the abiotic environment is identical in all samples (see supplemental methods section S1.4 for exceptions). A co-occurrence network was created from the *M* equilibrium abundances by computing the pairwise Pearson correlation between species abundances. A null distribution of Pearson correlations between abundances was created by simulating a network with no inter-species interactions, and correlations were considered significant if they fell outside the equal-tailed 99% confidence interval of that distribution.

### 2.2 Unipartite network construction

Random interaction networks were generated by first creating an Erdos-Renyi random graph [38]; the probability that an interaction coefficient *a*_*ij*_ is nonzero (i.e. the edge exists) is set to *p*_*ij*_. By default 100 species were used with *p*_*ij*_ = 0.05 and the probability of an interaction being positive set to *q* = 0.5.

Modular interaction networks were created by splitting 100 species into 5 modules of equal size, then assigning random interactions within each module. We then decreased the modularity of the network by re-assigning edges to random locations in the full network. This process will eventually converge to an Erdos-Renyi random graph with *p*_*ij*_ ≈ 0.17 for our default parameters. The initial probability of an interaction within a module being nonzero was 0.9 and the probability of an existing interaction being positive was *q* = 0.5.

Hub interaction networks have a bimodal degree distribution with a few high-degree nodes (hubs) and many low-degree nodes. Hubs are defined as nodes in the top 5% of the degree distribution. A hub network was created by splitting the network into modules as above and defining one species per module as a hub. The hub was then connected to 90% of the other nodes in the module. Then other edges were added in the network uniformly at random with probability *p*_*ij*_. Networks were then made progressively less bimodal by increasing *p*_*ij*_ slowly from 0 to 0.4. The probability of an interaction being positive was *q* = 0.1 (lower due to stability criteria; see supplement S1.3).

Networks with power law degree distributions were generated using the Barabasi-Albert random attachment algorithm [39] as implemented in NetworkX version 3.5 in Python [40]. Networks with exponential degree distributions were created by generating an set of degrees from an exponential distribution and then using NetworkX to generate a network with that specific set of degrees. Erdos-Renyi random graphs were also generated using NetworkX. In all cases, the expected degree was set to 10, which corresponds to *p*_*ij*_ ≈ 0.05 for the Erdos-Renyi random graphs.

### 2.3 Unipartite network properties and metrics

All metrics were assessed on an undirected, unweighted version of the networks to facilitate comparability between interaction and co-occurrence networks.

Node centrality in interaction and co-occurrence networks was assessed using NetworkX [40] (see supplement S1.5). Modules were detected using Clauset-Newman-Moore greedy modularity maximization [41] as implemented in NetworkX [40]. A pair of species was defined as correctly clustered if the two species were either in the same cluster for both the interaction and co-occurrence networks, or the two species were not clustered together in both the interaction and co-occurrence networks.

Degree distributions were fitted and evaluated using the R package poweRlaw [42, 43] (See supplemental methods). This assesses whether we can confidently conclude that the degree distribution is likely a power law [44]. Additionally in order to test whether we can reject the random network degree distribution expectation (binomial), we also performed Chi-squared goodness of fit tests on binned degree distributions (See supplemental methods).

### 2.4 Bipartite network construction

Bipartite interaction networks were generated by splitting *S* species into two classes *V*_1_ and *V*_2_ (either two classes of mutualists, such as plants and pollinators, or a class of exploitees and exploiters, referred to as hosts and parasites for ease) For mutualists *i* and *j*, both *a*_*ij*_ and *a*_*ji*_ *>* 0. For a host *k* and parasite *l, a*_*kl*_ *<* 0, while *a*_*lk*_ *>* 0. Interaction matrices were sorted such that species *i* ∈ *V*_1_ ∀*i* ≤ *S/*2, *i* ∈ *V*_2_ ∀*i > S/*2, ensuring inter-class interactions (mutualism or parasitism) were found on the upper-right and lower-left blocks of the interaction matrix. Note that either of these blocks alone reflect the biadjacency matrices commonly used to represent bipartite networks.

We constructed nested bipartite networks and modular bipartite networks whose inter-class interactions ranged from perfectly structured to randomly assigned, while controlling for overall edge density (see section S1.2 for details).

### 2.5 Bipartite nestedness and modularity

Only the inter-class interactions (the biadjacency matrix) of the interaction network were used when quantifying bipartite nestedness and modularity. This allows for focus on patterns that occur within a specified type of species interaction, without excluding other types of interactions from community dynamics as a whole [45]. NODF scores (Nestedness metric based on Overlap and Decreasing Fill) [46] were used to quantify nestedness of biadjacency matrices. Barber’s Modularity *Q*_*b*_, a bipartite version of the unipartite modularity introduced by Newman and Girvan [47, 48], was used to quantify modularity of the biadjacency matrices. In each simulation, *Q*_*b*_ reflects the modularity score of an optimal partition determined by the BRIM algorithm, implemented with the CDlib Python package ([49], default parameters). By determining optimal node partitions for an underlying interaction network and its resulting co-occurrence network, we assessed clustering correctness similarly to the unipartite case. Full details for both metrics can be found in section S1.5.

NODF and *Q*_*b*_ scores for a given interaction or co-occurrence biadjacency matrix were considered significant if they fell above (significantly anti-nested/modular, if below) the 95% confidence interval generated from NODF and *Q*_*b*_ scores of 1000 random matrices with the same edge density (see S1.5). For each type of bipartite network, we simulated 100 pairs of interaction/co-occurrence networks ranging from perfectly nested/modular to random. Each network was assigned a 1 if its relevant structural metric was determined to be statistically significant, or 0 otherwise. To quantify the degree of nestedness or modularity at which significant structure is lost, we performed a logistic regression on this binary encoding with the generalized linear model function in the Python statsmodels package [50], and calculated the resulting p = 0.5 decision boundary.

### 2.6 Code availability

All code is available at https://github.com/Fiona-MC/TheoryCoOccur-pub.

## 3 Results

### 3.1 Detecting direct interactions

When inter-specific interactions are assigned randomly and independently (Erdos-Renyi random graphs), we find that even with the highest tested interaction strength, the false discovery rate (proportion of significant correlations that are not direct interactions) is 0.3415. There is a strong correlation between interaction strength and Pearson correlation of abundances (Figure 1B), but there are many more non-interacting species pairs compared to interactions (Figure 1C), which causes many false discoveries. The area under the receiver operating characteristic (ROC) curve for the same simulation is 0.752, indicating a moderate ability to distinguish between interacting and non-interacting pairs using pairwise Pearson correlation of the species abundance data (Figure 1D). For lower interaction strengths, that ability is significantly diminished, reaching *AUC*_*ROC*_ = 0.572 for an interaction strength 10 times lower than the intra-species effect (*F*_*max*_ = 1/10, Figure 1D). This indicates that edges in a co-occurrence network often do not indicate that the species directly interact, regardless of the cutoff used to determine statistical significance.

**Figure 1:**
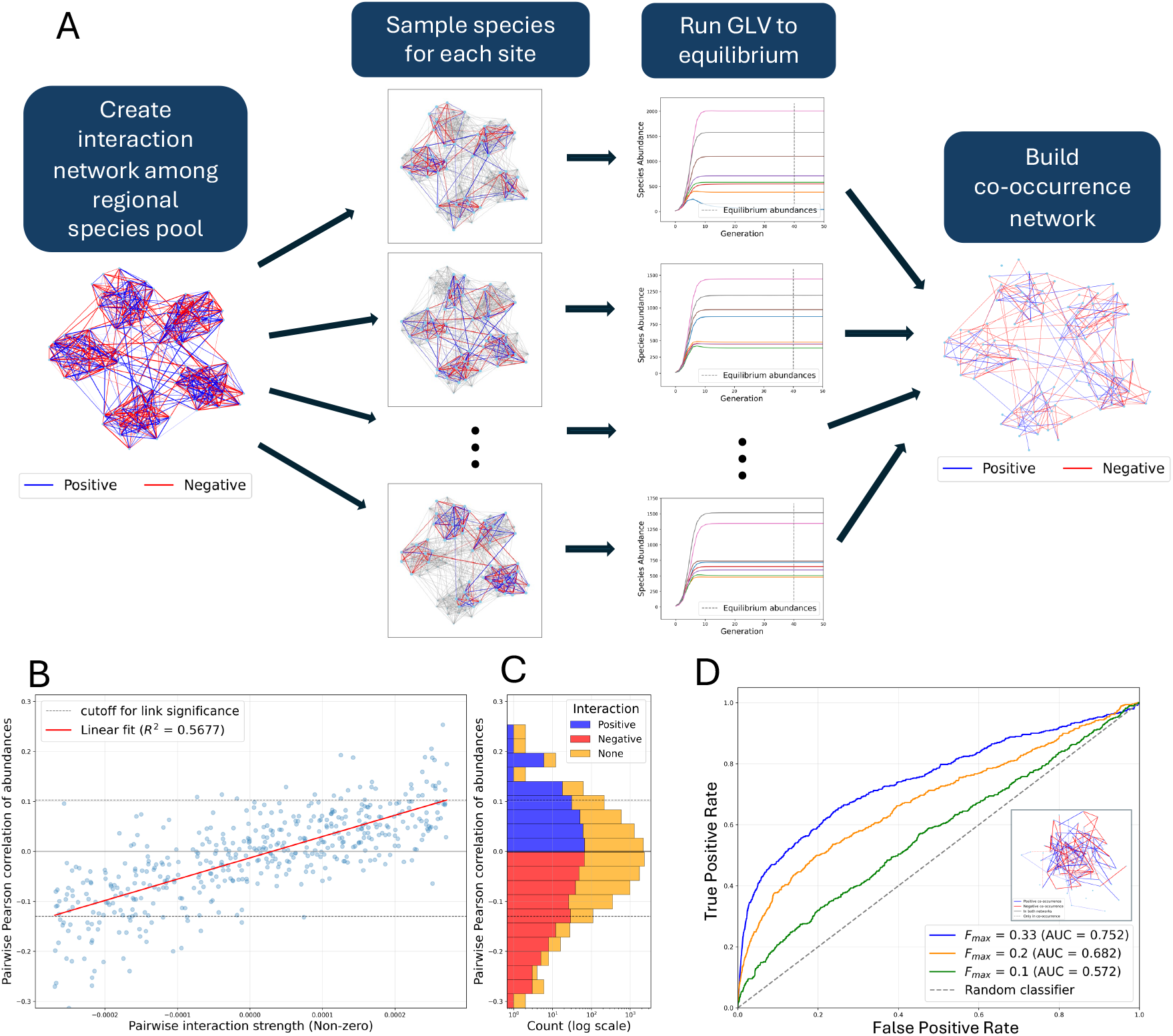
Many edges differ between the co-occurrence and interaction networks in simulated networks, in spite of a strong correlation between interaction strength and co-occurrence correlation. (A) Co-occurrence networks are generated from known interaction networks in a generalized Lotka-Volterra (GLV) based simulation. (B) Pairwise inter-specific interaction strength is highly correlated with pairwise co-occurrence relationships (*R*^2^=0.6099). Pearson correlation in pairwise species abundances versus interaction strength, and a linear fit to the data are shown for one Erdos-Renyi random interaction network. Non-interacting species pairs are excluded. Grey dotted lines show the 99% confidence interval of the null distribution of correlations. A co-occurrence edge is drawn between each species pair outside this interval. *F*_*max*_ = 1/5. (C) Distribution of pairwise Pearson correlations of species abundances for interacting and non-interacting species (log scale) for the same simulation as in B. Blue indicates a positive interaction between the species, red indicates a negative interaction, and yellow indicates no interaction. (D) Receiver operating characteristic (ROC) curves for direct edge prediction compared across three interaction strengths (*F*_*max*_ ∈ {1/3, 1/5, 1/10}) in an Erdos-Renyi random graph. True positives are defined as undirected edges that exist in both the interaction and co-occurrence network. Inset network shows one example of a co-occurrence network, where the dotted lines represent edges that are in the co-occurrence network but not in the interaction network. Area under each ROC curve (AUC) is shown in the legend.

We also measured how well the co-occurrence network preserves the sign of the interaction (positive versus negative). We see that the proportion of positive interactions is strongly correlated with the proportion of significant positive correlations in the co-occurrence network (Supplemental Figure S1). However, while the interaction network proportion of positive edges ranges from 0 to 0.8, the co-occurrence proportion only ranges from 0.4-0.55 (Supplemental Figure S1). We were not able to create networks with greater than 80% positive interactions due to GLV system instability (See supplement S1.3).

### 3.2 Hub species and node centrality

Although many edges in the co-occurrence network do not represent direct interactions (Figure 2B), we find that we can predict high-degree nodes in the interaction network (hubs) using metrics of centrality on the co-occurrence network (Figure 2A). Here, we test the ability to distinguish between high and low degree nodes in networks with highly defined hubs, and then add interactions throughout the network to gradually erode the difference between the hubs and the other nodes. Here, *AUC*_*ROC*_ measures the probability that a randomly chosen hub node will have a higher centrality metric than a randomly chosen non-hub node. When the hub species interact with almost all of the species in their module (on average 90%), and there are no other interactions, the average area under the ROC curve was 0.999 for degree centrality. Betweenness centrality performs nearly as well for detecting high-degree nodes (mean *AUC*_*ROC*_ = 0.987), whereas closeness and eigenvector centrality do not perform as well (*AUC*_*ROC*_ = 0.905 and *AUC*_*ROC*_ = 0.917 respectively; Figure 2A). It is not surprising that degree centrality would perform well given that the definition of a hub here is that the node has high degree in the network. The lower performance of closeness centrality in this case may be due to the modular structure of the graph, since it is expected that hub nodes will be far from other modules.

**Figure 2:**
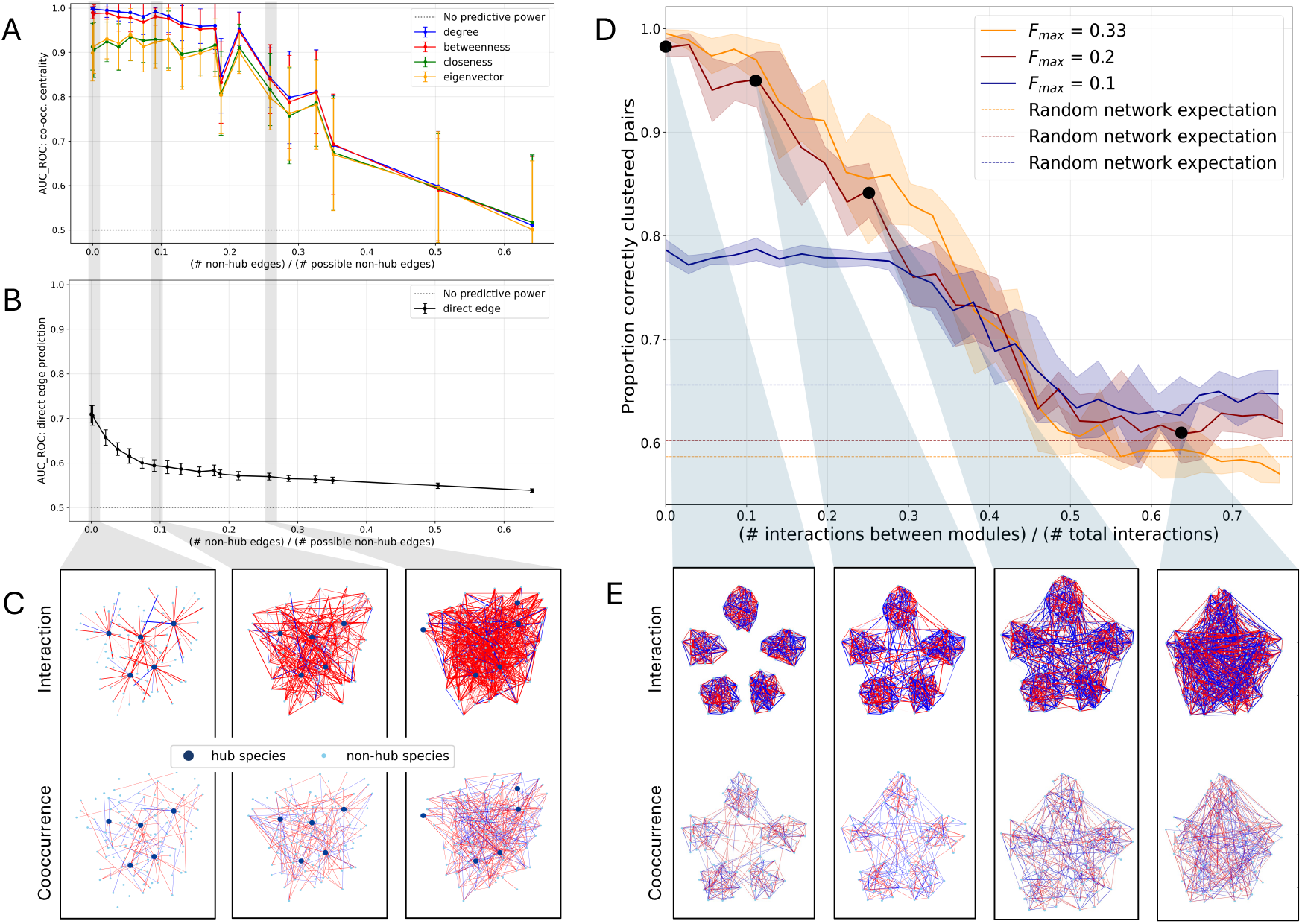
High degree nodes (hubs) and modules in the interaction network can be predicted from the co-occurrence network in many cases, in spite of many differences in direct edges. (A) Interaction network hub detection (*AUC*_*ROC*_: co-occ. centrality) for different amounts of hub degree differentiation ((# non-hub edges)/(# possible non-hub edges)). The *AUC*_*R*_*OC* value is computed for prediction of the top 5 highest degree nodes in the interaction network using four metrics of co-occurrence node centrality. Points are are averages over 30 replicates and error bars are the standard deviation of the replicates. *F*_*max*_ = 1/5 and *q* = 0.1. Edge proportions - (# non-hub edges)/(# possible non-hub edges) - are computed for the undirected interaction network. (B) *AUC*_*ROC*_ for direct interaction edge prediction using Pearson correlation in the co-occurrence data, computed on the same networks as in A. (C) Examples of interaction (top) and co-occurrence (bottom) networks with five highest degree nodes marked as hubs (larger blue nodes). Red indicates negative interaction/co-occurrence, blue indicates positive, and the edge width indicates interaction/correlation strength. (D) Proportion of pairs of nodes that are correctly clustered versus the degree of modularity (proportion of the realized edges in the interaction network that are between modules). A pair of nodes is defined as “correctly clustered” if both nodes are in the same cluster in the interaction network and in the co-occurrence network, or if the two nodes are in different clusters in both the interaction and co-occurrence networks. Realized edge proportion - (# interactions between modules)/(# total interactions) - is computed for the undirected interaction network. Three lines are shown for interaction strength factor (*F*_*max*_) of 1/3, 1/5, and 1/10. Error bars are shown for the standard deviation of five replicates. Random network expectations are shown for the same metric computed for an Erdos-Renyi random graph with the same number of edges and the same maximum interaction strengths. (E) Examples of interaction and co-occurrence networks at different levels of module deterioration. All interaction networks have the same total number of edges. Co-occurrence edges, edge colors, and edge widths are defined the same as in C. *F*_*max*_ = 1/5 and *q* = 0.5.

As edges are added to the baseline hub network, bringing the degree of non-hub nodes closer to that of the hubs, centrality measures on the co-occurrence network remain effective predictors of high node degree in the interaction network. When the interaction network has nearly 20% of the edges not connected to hub nodes realized, the *AUC*_*ROC*_ for predicting the top 5 highest degree nodes using co-occurrence node degree centrality is still over 90% (Figure 2A).

For the same networks, we are not able to predict the interaction edges well using Pearson correlation of the co-occurrence data (Figure 2B). This indicates that although a high percentage of the edges in the co-occurrence network do not represent direct interactions, nodes that had high degree in the interaction network still have high degree in the co-occurrence network.

### 3.3 Detecting unipartite modules and modularity

Identifying modules of species that interact more closely with one another than with other community members is often of interest in ecological contexts. Therefore, we explored the ability of co-occurrence networks to retain signatures of modules within interaction networks. When comparing the node groupings identified by a clustering algorithm applied to both an interaction network and its resulting co-occurrence network, we find that we can identify similar modules for both networks over a wide range of interaction network modularity (Figure 2D).

For higher interaction strengths, the clustering algorithm clusters nodes in the co-occurrence network almost identically to the interaction network when the interaction network modules are disconnected. Even for low interaction strengths, co-occurrence networks generated from highly modular interaction networks correctly cluster about 80% of nodes. As we move interactions from inside the module to random locations in the network, the ability to detect the same modules deteriorates. However, until approximately 40% of the edges in the interaction network are between modules, the clustering correctness of co-occurrence networks exceeds the random network expectation.

### 3.4 Differentiating between node degree distributions

If network edges are assigned randomly and independently (Erdos-Renyi random graphs), a binomial degree distribution is expected. Real-world networks in a variety of fields have a heavier tail than expected under this model, and in many cases these distributions have been characterized as power laws [25, 51, 52]. However, it is very difficult to distinguish between power laws and other heavy-tailed distributions, such as the log-normal distribution, particularly in networks with small numbers of nodes like those seen in community ecology [44, 53]. Using statistically robust methods to differentiate between distributions [44], over 10^5^ nodes (i.e. species) are needed to differentiate between power law and log-normal distributions even under perfect conditions (Table S2.1). Therefore, even when the interaction networks are generated to have power law degree distributions, we cannot detect this robustly when comparing to other heavy tailed distributions. We can see that for networks that are generated from distinct node degree distributions, multiple distributions may fit the data similarly well (Figure 3B, Table S2.2). In our set of tests, we were never able to confidently conclude that any node degree distribution was a power law (Table S2.2).

**Figure 3:**
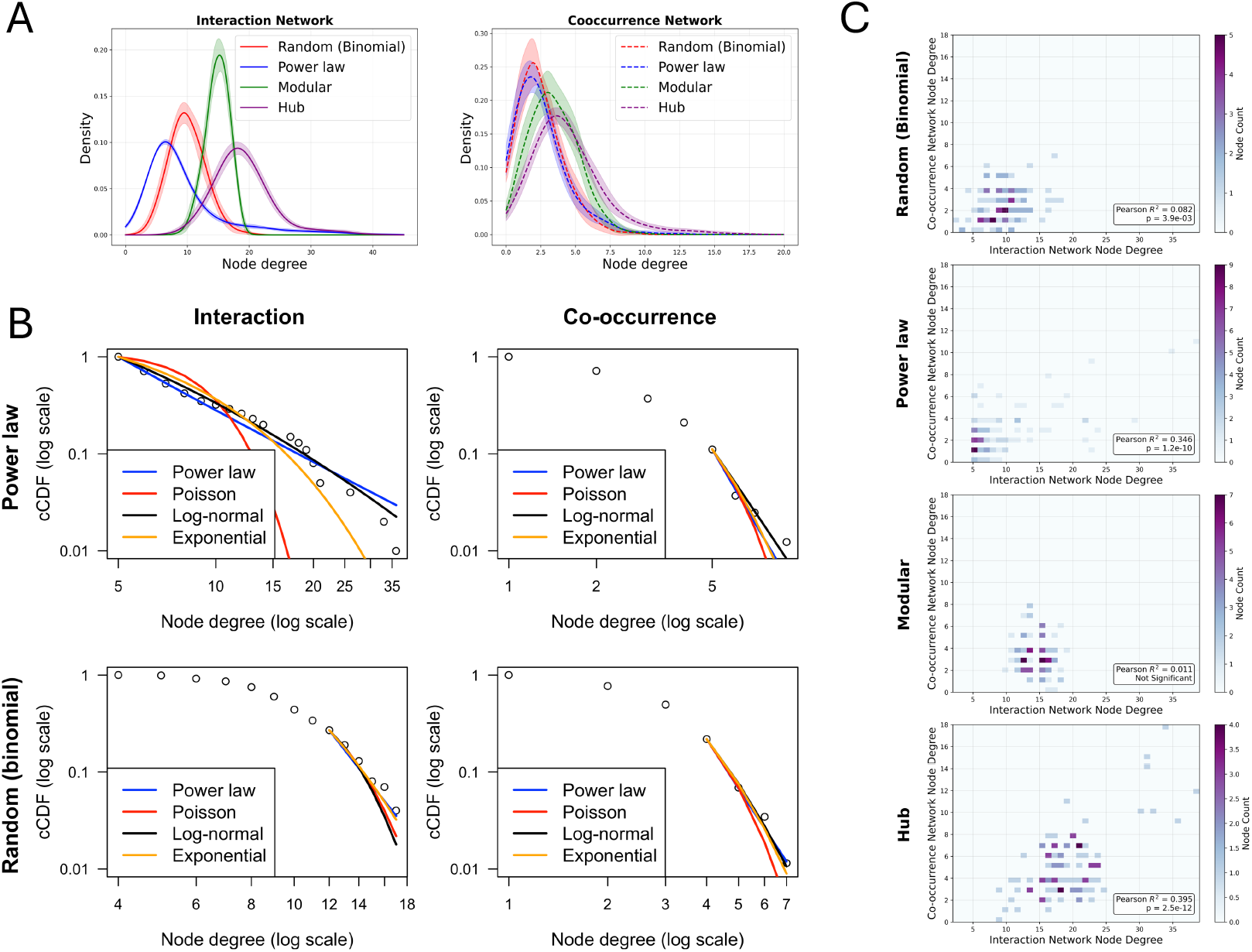
Family of degree distribution cannot be reliably detected for interaction or co-occurrence networks. (A) Degree distributions (estimated using kernel density estimation) for the interaction and co-occurrence in networks with 100 taxa (nodes). Lines are the average over 10 networks, and shaded interval is one standard deviation of the replicates. *F*_*max*_ = 1/5. (B) Empirical complementary cumulative distribution function (cCDF) of the degree distribution (open circles), and maximum likelihood fits to the empirical degree distributions for four types of distributions (colored lines). Results are shown for the interaction and co-occurrence networks with 100 taxa (nodes). The true node degree distribution of the interaction network is shown on the left. All fits use data only for degrees greater than *x*_*min*_ which is fitted for the power law distribution. Note: Poisson distribution is a reasonable approximation of the binomial distribution with these parameters. (c) Degree comparison per node between interaction and co-occurrence networks for interaction networks with differing degree distributions. Network with 100 taxa (nodes) and interaction strength factor *F*_*max*_ = 1/5. Color indicates the number of nodes at each point on the graph. Pearson *R*^2^ and p-value for a linear fit is shown on each plot.

However, we hypothesized that it may nonetheless be possible to conclude that the distribution of node degrees differs significantly from a binomial distribution, and that this property may be retained from the interaction network to the co-occurrence network. As expected, when tested on the ground truth interaction network node degree distributions, we incorrectly reject the null hypothesis 4% of the time for Erdos-Renyi random graphs. We also correctly reject the null hypothesis for 100% of power law and exponential interaction networks. For the co-occurrence network degree distributions, we reject the binomial hypothesis more often when the interaction network had a power law degree distribution compared to when the interaction network had a binomial degree distribution. However, this result was sensitive to interaction strength and number of species in the network (Table S2.3). This suggests that the co-occurrence network does not necessarily have a binomial degree distribution, even when the interaction network does. Correspondingly, the degree distributions of co-occurrence networks are qualitatively more similar to one another than their generative interaction networks (Figure 3A).

Though the interaction network degree distributions are not preserved, the degrees of the individual nodes in the interaction network are correlated with the degrees of the nodes in the co-occurrence network across multiple interaction node degree distributions (Figure 3C). Correlations are strongest for hub interaction networks, and lowest for modular interaction networks.

### 3.5 Detecting bipartite nestedness and modularity

Many ecological network studies center on bipartite networks. In bipartite networks, species are divided into two classes that engage in a single type of inter-class interaction. For both bipartite mutualist and bipartite host-parasite assemblages, we examine the degree to which common metrics of nestedness and modularity are retained in co-occurrence networks. Broadly, nestedness describes the degree to which specialist interactions are subsets of generalist interactions, while modularity describes the degree to which a network is compartmentalized into subsets that interact more regularly with one another than with other subsets. We construct interaction networks in which inter-class interactions range from perfectly nested or modular to random, and assess the structural features of their corresponding co-occurrence networks. In our simulations, a decline in probability of a nested or modular interaction is balanced by an increase in the probability of a non-nested or non-modular interaction, maintaining the same expected interaction density across the biadjacency matrix (see S1.5.2).

#### 3.5.1 Nested Bipartite Networks

Nestedness is significantly detected up to a 43% probability of a non-nested interaction in underlying interaction networks (p = 0.03, pseudo-*R*^2^ = 0.390), and up to a 24% probability of a non-nested interaction in co-occurrence networks (p *<*0.001, pseudo-*R*^2^ = 0.539) (Figure 4). Thus, co-occurrence networks of nested mutualists remain significantly nested across most of the degrees of nestedness tested, though they fail to retain this signal as long as their generative interaction networks.

**Figure 4:**
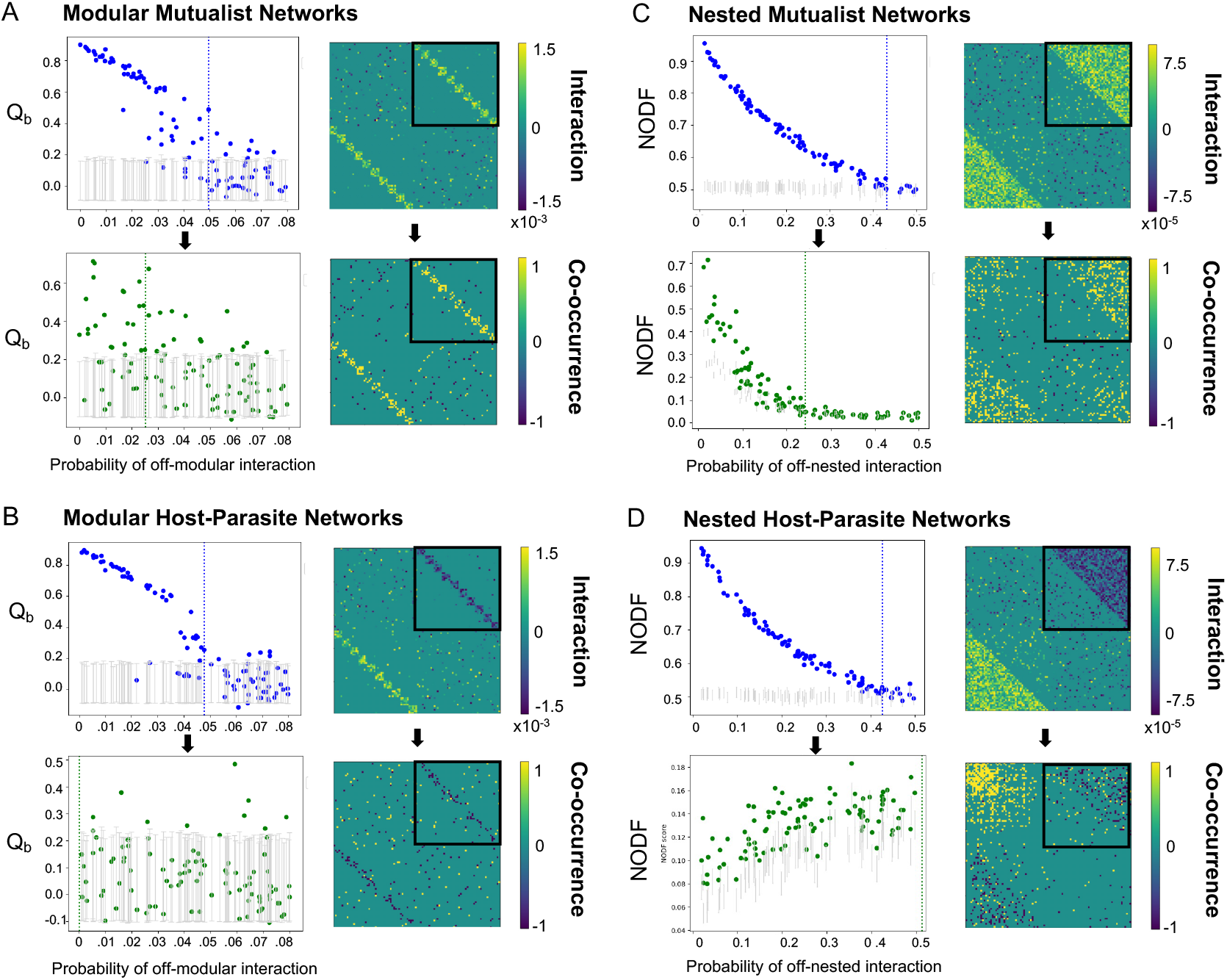
Detecting modular and nested structures in bipartite interaction and co-occurrence networks of modular mutualist (A), modular host-parasite(B), nested mutualist (C), and nested host-parasite (D) networks. **(A, B)** 100 interaction networks with varying degrees of modularity (i.e. probabilities of an off-modular interaction) were simulated, as described in section S1.2. Intra-class interactions were assigned with probability *p*_*ij*_ = 0.05 and *q* = 0.5. *Q*_*b*_ scores were calculated for the biadjacency matrix (boxed in black in the representative adjacency matrices located at right, for *P* (off-modular interaction) = 0.01) of both the interaction network and corresponding co-occurrence network. **(C, D)** 100 interaction networks with varying degrees of nestedness were simulated. Intra-class interactions were assigned with probability *p*_*ij*_ = 0.1 and *q* = 0.2. NODF scores were calculated for the biadjacency matrix (boxed in black in the representative adjacency matrices located at right, for *P* (off-nested interaction) = 0.1) of both the interaction network and corresponding co-occurrence network. **(A-D)** Grey bars represent 95% confidence intervals for the respective network metric, based on 10^3^ random permutations of the interaction or co-occurrence network. Vertical dotted lines indicate a threshold beyond which we fail to detect significant nestedness or modularity more than half the time, modeled using logistic regression of a binary encoding of the presence of statistically significant nestedness or modularity.

Nested host-parasite networks do not follow the same trends as their mutualist counterparts. Lower degrees of nestedness in host-parasite networks leads to both higher edge density and *higher* NODF (Nestedness metric based on Overlap and Decreasing Fill) scores in resulting co-occurrence networks, and these co-occurrence networks tend to be significantly nested even after the interaction networks become statistically indistinguishable from a random bipartite network (up to a 42% probability of a non-nested interaction; p = 0.023, pseudo-*R*^2^ = 0.4883) (Figure 4). Thus, a signal of significant nestedness is retained, but not a directly informative one. When an alternative, partially constrained null model was employed – in which entries are assigned probabilistically such that the expected marginal totals are held constant [54] (see S1.5.2 and Figure S3C) – co-occurrence networks no longer exhibit significant nestedness. Intriguingly, this implies that in the case of host-parasite co-occurrence networks, nestedness was more strongly driven by heterogeneity in co-occurrence degree distribution (which emerged even in largely random consortia) rather than proper subsetting of specialist interactions within generalist interactions.

#### 3.5.2 Modular Bipartite Networks

For modular mutualist networks, *Q*_*b*_ modularity scores in both interaction networks and co-occurrence networks exhibited a similar decay as the probability of an off-module interaction increased. Like the nested mutualist networks, the modular mutualist co-occurrence networks remained significantly modular at higher degrees of modularity, but did not retain this signal as long as their generative interaction networks across the tested degrees of modularity. Modularity is significantly detected in interaction networks with up to a 4.9% probability of an off-module interaction (p *<*0.001, pseudo-*R*^2^ = 0.4272), and up to a 2.5% probability of an off-module interaction in co-occurrence networks (p *<*0.001, pseudo-*R*^2^ = 0.1778) (Figure 4). Note the perfect modular bipartite construction has a lower overall edge density than the perfect nested construction, leading to the lower thresholds for retaining significant structure. Figure 4 shows results from a 10-module structure; when species are grouped instead into 5 looser modules (see Figure S3), co-occurrence network *Q*_*b*_ scores do not remain significantly modular at any degree of interaction network modularity. For modular host-parasite networks, *Q*_*b*_ scores of interaction networks and co-occurrence networks showed little correlation; co-occurrence networks were not significantly modular even with nearly perfect modular interaction networks.

Modularity scores in each simulation are based on a parition of nodes from an independent application of the BRIM algorithm, not a fixed partition. To determine if the species composition of each module remains consistent, we compared cluster membership in the partitioned co-occurrence network to the cluster membership of the interaction network. For highly modular mutualists, clustering correctness can exceed 80-90% (Figure 5). This is partially driven by the disproportionate numbers of species pairs (correctly) not clustered together (i.e., captures true negatives; this is why even clustering correctness of a random bipartite network is quite high). When considering only true positives (pairs of species clustered together in the co-occurrence network that also clustered together in the interaction network), co-occurrence networks of highly modular host-parasite bipartite networks correctly identified 20-40% of clustered pairs. Co-occurrence networks of highly modular mutualist bipartite networks correctly identify *>*75 % of clustered pairs. Therefore, although highly modular host-parasite interaction networks (and less-densely interacting modules of mutualists) did not generate co-occurrence networks with signals of significant modularity beyond that of a random interaction network, identified modules were more accurate than would be expected from an underlying random network.

**Figure 5:**
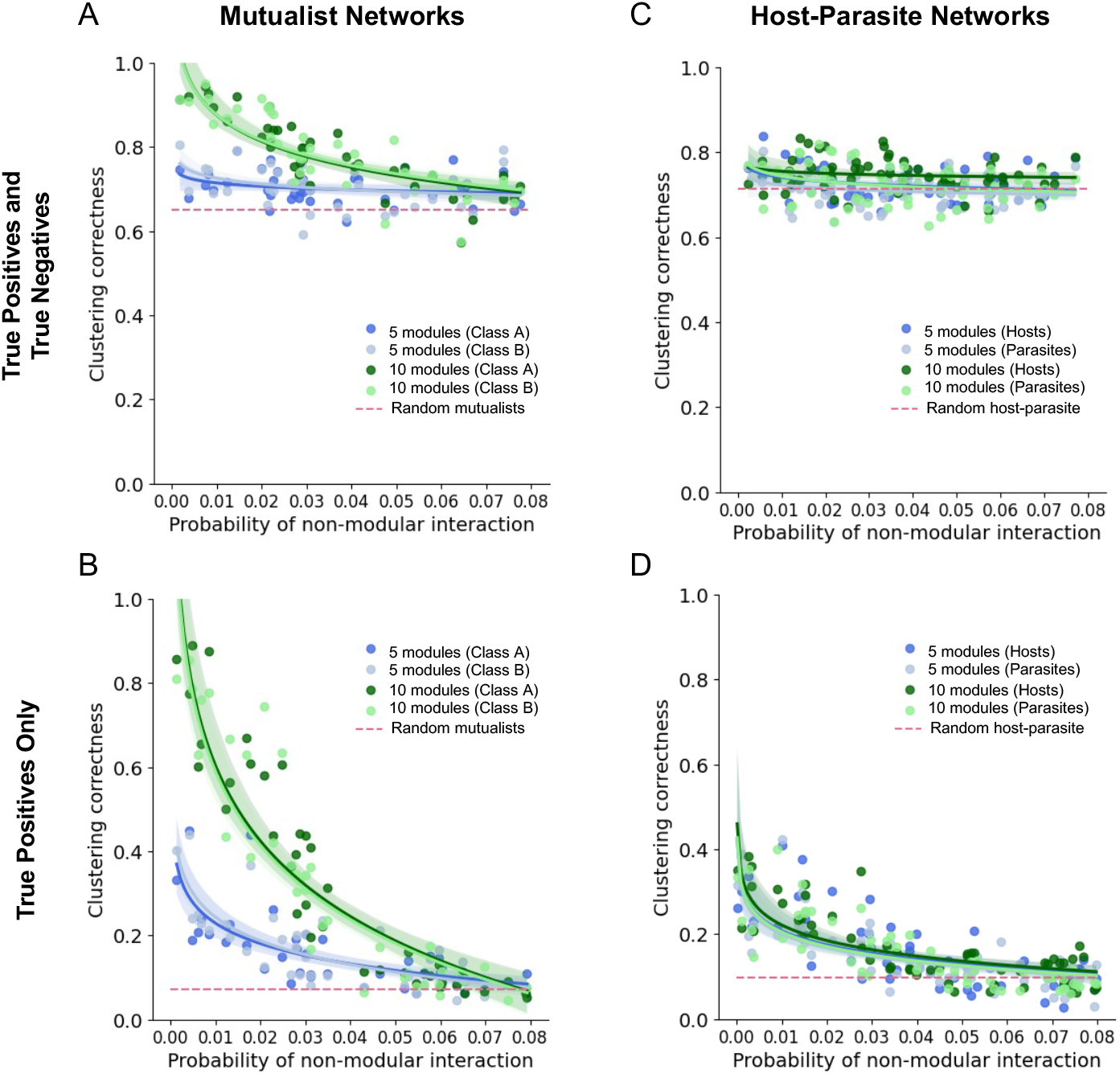
Identifying clusters of interacting species in modular mutualist **(A, B)** and host-parasite **(C, D)** bipartite networks. 100 interaction networks with varying degrees of modularity were simulated over the same range and with the same parameters as in Figure 4C-D, with interactions split either into 5 or 10 modules. Bipartite clustering was performed with BRIM algorithm for both the interaction network and resulting co-occurrence network. Total pairwise clustering accuracy of the co-occurrence network relative to the node partitions identified for the interaction network are shown in (A) and (C), plotted by class. Successful detection of the existing clusters in the interaction network (i.e. true positives only) are shown in and (D). Shaded areas represent a 95% CI for the linear-log regression. Horizontal dashed lines represent the pairwise clustering accuracy obtained when performing bipartite clustering on a bipartite network with random inter-class interactions and its resulting co-occurrence network.

## 4 Discussion

A compelling reason to develop network representations of ecological communities is to better understand the broader network motifs - beyond pairwise species interactions - that influence community-level processes. The issues with using co-occurrence data to infer direct, pairwise species interactions are well documented [19, 24]. Patterns of species co-occurrence are emergent consequences of the aggregate effect of many interactions; we therefore hypothesized that co-occurrence networks, while failing to accurately reflect pairwise interactions, may nevertheless reflect emergent network structures hypothesized to drive collective processes in ecology.

### 4.1 Co-occurrence networks can retain emergent interaction structures

It is known that under real-world conditions, many correlations in co-occurrence networks are caused by indirect interactions (either through shared responses to the environment or interactions through other species), non-independence in time and space, and other violations of modeling assumptions [19, 24]. In this idealized simulation setting, external environmental factors are removed, communities are at equilibrium, and all data points are independent, allowing us to disregard these important aspects of the source of correlation between species in real data. However, even in this best-case scenario, we still observe a large proportion of significant correlations that do not represent direct interactions.

Nevertheless, we find that co-occurrence networks can reproduce patterns seen in the corresponding interaction networks, despite a high percentage of edges differing between the networks. By systematically investigating the detection of hub species, unipartite modularity, degree distributions, and bipartite nestedness and modularity, we find that the very features of a network that give rise to higher order structures – heterogeneity in interaction densities and biased edge attachment – are qualitatively retained in co-occurrence data.

Heterogeneities in interaction density are reflected in co-occurrence data, as seen in successes in detecting hub species (species with high node degree), and in the broadly generalizable positive correlations between interaction node degrees and co-occurrence node degrees. This suggests that we may have success in identifying more cosmopolitan species compared to specialist species from co-occurrence data, narrowing the scope of future lab or field experiments required to identify the particular direct interactions of interest. In contrived interaction networks with narrow (low heterogeneity) degree distributions (e.g. the unipartite modular networks, where species had far more similar numbers of interactions than would be expected randomly), we observe a lower correlation between interaction and co-occurrence node degrees (Figure 3C).

The retention of heterogeneous interaction densities is also reflected by the retention of nestedness in bipartite co-occurrence networks. Highly nested bipartite interaction networks have heterogeneous degree sequences; indeed, recent theoretical work highlights how nestedness can be an entropic consequence of heterogeneous degree sequences in a network [55]. For bipartite mutualist networks, this signal of nestedness was retained in co-occurrence networks over a wide range of interaction network nestedness.

In addition to heterogeneous interaction densities, biased edge attachment in interaction networks are observed in co-occurrence networks. In particular, both unipartite and bipartite modular structures in interaction networks can persist in co-occurrence networks. We hypothesize that dense modules self-reinforce through indirect chains of interactions, meaning module identity and overall degree of modularity are retained (Figure S4). The greater degree of clustering accuracy in modular mutualist networks relative to modular host-parasite networks highlights the importance of this self-reinforcement, as dense clusters of mutualists experience strong positive feedback that translates to strong positive correlations among species abundances.

### 4.2 Ecological network degree distributions remain elusive

There is long-standing interest in determining if heavy-tailed degree distributions, which are often seen in other complex systems, are seen in ecological contexts. We find that it is very difficult to confidently distinguish between candidate distributions using rigorous statistical tests [44]. Even with the ground truth interaction network degree distribution, we were unable to confidently classify the degree distributions with networks that have reasonable sizes for ecological communities.

Previous work has suggested that biotic bipartite interaction networks follow power laws and co-occurrence networks follow exponential distributions [25]. However, it is likely that they cannot confidently distinguish between power laws and other distributions using rigorous statistical tests. Several methods have been used to draw conclusions about networks following power laws. One is to compare the fits of different distributions and take the best fit to be the correct distribution. This type of test will often come to the incorrect conclusion because it does not rigorously compare the fit of different distributions (for example using a p-value from a likelihood ratio-based test) [25, 44]. Additionally, it is common to use the p-value from a linear fit between the log of the empirical CDF of the degree distribution and the log of the node degrees. This tests the hypothesis that these values are significantly correlated. However, this does not imply that the linear fit is the best functional form, and therefore is not a statistically rigorous way to test whether a distribution follows a power law [44].

### 4.3 Looking towards empirical data: reframing co-occurrence networks as tools for exploring hypotheses

To create co-occurrence networks, we compute Pearson correlations between abundances of pairs of species produced by a GLV simulation. In empirical studies, more complex methods are used to account for various factors (e.g. accounting for compositional data in metagenomics data), and some methods have been employed to attempt to account for indirect interactions through other species and the environment [22, 23, 56–58]. However, previous work has indicated that many edges in these co-occurrence networks still do not represent direct interactions [24]. There is additional promise in dynamic Bayesian networks, which aim to infer causality by using information from time series, but large amounts of very high-resolution time series data are needed for reliable inference [59].

Furthermore, environmental drivers are important factors in real-world ecological networks. It is well established that in empirical studies, co-occurrence relationships are caused by shared responses to the abiotic environment or to unmeasured biotic factors. For example, a module in the co-occurrence network may comprise a set of species that respond similarly to a shift in the abiotic environment, reflected by the greater similarity in species’ environmental preferences within a module than between modules (Fig. S5). It is thus important to consider multiple alternative hypotheses about the observed network patterns, in the context of known environmental variation.

We want to be intentional in framing our work not as a guarantee of successful inference when applied to real biological data. Rather, we propose an avenue for moving from collecting co-occurrence data to gaining insights into ecological communities – without requiring a definitive pairwise interaction network.

There still exists a large gap between the theory hypothesizing that emergent network structures play key roles in community function, and lab or field experiments demonstrating that these structures are important in real-world settings. Indeed, the assumptions undergirding influential hypotheses from ecological theory, such as the stabilizing role of nestedness in bipartite interaction networks [60], or the modular structure expected from a coevolutionary system with a matching allele model [61], are challenged when tested in empirical systems [45, 62]. If the outputs of ecological network analysis are to extend beyond descriptive analysis [63], experimental tests of hypotheses from ecological network theory are critical.

Our work suggests that many of the proposed influential features of ecological interaction networks – modules, hierarchical interaction structures, sets of super-generalists – may be retained in co–occurrence networks. Thus, rather than focusing on the identity of individual pairwise interactions, these groups and features identified in co-occurrence networks could be used in future lab and field experiments studying the role of broader network topology in shaping outcomes of species removal, species invasions, community coalescence, environmental perturbations, and coevolution. Recent work linking changes in co-occurrence network topology to functional outcomes in plant microbiomes suggests this is a rich area for study [64], and we hope to motivate future work utilizing co-occurrence data to better connect hypotheses from network theory with ecological community function.

## Notes

### Competing Interest Statement

The authors have declared no competing interest.

### Funding

- National Science Foundation: NSF-DGE-2146752

