## Supplemental materials for "Co-occurrence networks can preserve emergent properties of ecological communities"

### S1 Supplementary Methods

| Variable | Definition | Default setting |
|---|---|---|
| $t$ | Time index | - |
| $M$ | Number of samples | 500 |
| $S$ | Number of nodes (species) | 100 |
| $S_m$ | Number of nodes subsampled | $S/2$ |
| $p^{(rand)}$ | Proportion of interactions in random network | 0.05 |
| $p^{(mod)}$ | Initial proportion of interaction within a module for the unipartite modular networks | 0.9 |
| $p^{(hub)}$ | Initial proportion of interaction within a module from a hub node for hub networks | 0.9 |
| $q$ | Proportion of realized interactions that are positive | 0.5 |
| $I_{max}$ | Max interaction strength | $\alpha \cdot F_{max}$ |
| $F_{max}$ | Max ratio between the inter-species and intra-species effect strength | 1/5 (also 1/3 and 1/10 for select analyses) |
| $\boldsymbol{n}(t)$ | Species abundances | $n_i(0) \sim Uniform(10, 20)$ |
| $\boldsymbol{b}$ | Intrinsic growth rates | $b_i \sim Uniform(0.9, 1)$ |
| $\alpha$ | Self-limitation parameter for all species (varies only in environment simulations; see Methods) | 0.001 |
| $\boldsymbol{A}$ | Community matrix of Lotka-Volterra interaction parameters | See section on interaction network generation. |
| $E$ | Environment value (For environment simulations only) | $E \sim Bernoulli(0.5)$ |
| $\tau_i$ | Environmental preference for species $i$ (only for environment simulations) | $\tau_i \sim Unif(0, 1)$ |

Table S1: Parameters and values used in standard simulations. Note $\alpha$ is kept constant for simplicity, as previous work [1] demonstrated the relationship between an interaction network and co-occurrence network is robust to species evenness (when sampling rates are not a concern). This is verified for our system in section S4.

#### S1.1 Generating unipartite interaction networks

For the analysis where we made it harder and harder to detect the hubs, we made five modules, each with a hub that is fully connected to the other species in the module, and then made it harder and harder to detect these hubs by increasing the probability of an interaction elsewhere in the network. Hubs were defined as the nodes in the top 5% of the degree distribution. This means that eventually as we increase $p_{ij}$, some other nodes by chance might become hubs, pushing some of the original hubs out of the top 5%. One hundred independent networks were averaged for each level of $p_{ij}$ in the results for this analysis. $p_{ij}$ ranged from 0 to 0.2.

The probability of a positive interaction (given any interaction exists) for these networks was set to 0.1 to avoid instability in the system (See Section S1.3).

#### S1.2 Generating bipartite interaction networks

Bipartite interaction networks were generated by splitting $S$ species into two classes $V_1$ and $V_2$ (either two classes of mutualists, such as plants and pollinators, or a class of exploitees and exploiters (referred to as hosts and parasites for ease). Interaction matrices were sorted such that species $i \in V_1$ $\forall i \leq S/2$, $i \in V_2$ $\forall i > S/2$, ensuring inter-class interactions (mutualism or parasitism) were found on the upper-right and lower-left blocks of the interaction matrix. For nested bipartite networks (which had a higher density of interactions), intra-class interactions existed with probability $p_{ij} = 0.1$ for $i, j \in V_1$ and $i, j \in V_2$, with probability $q = 0.2$ of a positive interaction (avoiding system instabilities driven by an excess of positive interactions). For modular bipartite networks, intra-class interactions existed with probability $p_{ij} = 0.05$ for $i, j \in V_1$ and $i, j \in V_2$, with probability $q = 0.5$ of a positive interaction.

Perfectly nested bipartite networks had inter-class interaction blocks that contain upper and lower triangular matrices. To systematically reduce the degree of nestedness, the probability of an interaction was reduced for edges falling within ($p^{(w)}$) the the upper (lower) triangular structure and increased for edges falling below (above) the main diagonal ($p^{(o)}$). To keep expected total number of inter-class interactions constant, we ensure $p^{(w)} = 1 - p^{(o)}$, due to symmetry across the block diagonal. For perfect nestedness, $p^{(w)} = 1$ and $p^{(o)} = 0$. We vary $p^{(w)} \in [0.5, 1]$, ranging from 0 (perfect nestedness) to 0.5 (random interactions).

To vary the degree of modularity within bipartite interaction structures, inter-class interaction network edges were created with probability $p^{(m)}$ for interactions occurring within a module, and probability $p^{(n)}$ for interactions occurring outside a module. To keep the expected total number of inter-class interactions constant, $p^{(m)}$ ($p^{(n)}$) was decreased (increased) in a module-size dependent manner, determined as follows:

For a module of size $m$, $m$ species of class $V_1$ interact with $m$ species of class $V_2$. Therefore, for $S$ total species, the number of possible within-module edges in either the upper right or lower left block of the interaction matrix $|L_m|$ can be found:

$$|L_m| = \frac{S}{2m}m^2 = \frac{Sm}{2}. \tag{S1}$$

The total number of possible non-modular edges in either the upper right or lower left block of the interaction matrix then follows as

$$|L_n| = \frac{S^2}{4} - |L_m| = \frac{S^2}{4} - \frac{Sm}{2} = \frac{S}{4}(S - 2m). \tag{S2}$$

We therefore ensure that

$$-\Delta p^{(m)}|L_m| = \Delta p^{(n)}|L_n|, \tag{S3}$$

or

$$-\Delta p^{(m)} = \Delta p^{(n)}\frac{S - 2m}{2m}. \tag{S4}$$

#### S1.3 Stable equilibria for community simulations

When simulating the GLV system for each subsampled community, equilibrium was assessed by requiring that the abundances change less than 1% of between time points. In some parameter and structural combinations, positive interactions between species cause the system to enter a positive feedback loop where convergence does not occur. This is a known property of GLV models; mechanisms of avoiding runaway positive feedback have been reviewed previously, with appropriate model choices depending on ecological context [2–5]. Rather than arbitrarily selecting a more complex functional form for our generalized species interactions, we note clearly in the text where biologically infeasible cases were avoided by either a) lowering the maximum strength of interactions by lowering $F_{max}$, b) lowering the proportion of interactions ($p_{ij}$) or c) lowering the proportion of positive interactions ($q$).

#### S1.4 Simulating co-abundance from environmental preferences

For most simulations in the paper, species do not respond to an abiotic environment. However, in a select few simulations, we added this factor to explore how network structures can change in a non-homogenous environment. To model variable environmental preferences among species, we used a uniform distribution to assign a trait to each species, and then compared the trait to the current simulated environment. The carrying capacity of the species was adjusted based on the mismatch between the preference and the environment.

In an ideal environment, $\alpha$ is the self-limitation parameter; $\alpha_i$ is the self-limitation parameter for for species $i$ after adjusting for the trait-environment mismatch. Let $E \in \{0, 1\}$ be the value of the environment and $\tau_i \sim Unif(0, 1)$ be the trait value for that species. We then assign

$$\alpha_i = \frac{\alpha}{1 - 0.5 \cdot |E - \tau_i|} \tag{S5}$$

Environments are either set to all be 0 (null) or set to $E \sim Bernoulli(0.5)$ for each data point. This corresponds to reducing the base carrying capacity (in the absence of interspecific interactions) by a percentage defined by the environment mismatch.

#### S1.5 Network properties and metrics

##### S1.5.1 Node centrality

Node centrality in interaction and co-occurrence networks was assessed by several metrics: degree centrality, betweenness centrality, closeness centrality, and eigenvector centrality. All metrics were implemented using functions in NetworkX [6]. Node centrality was assessed on an undirected, unweighted version of the networks to facilitate comparability between interaction and co-occurrence networks.

##### S1.5.2 Bipartite nestedness and modularity

Only the inter-class interactions (the biadjacency matrix) of the interaction network were used when quantifying bipartite nestedness and modularity. Entries in the biadjacency matrix were converted to 1s and 0s to reflect the presence or absence of an interaction, respectively.

**Nestedness:** NODF scores (Nestedness metric based on Overlap and Decreasing Fill)[7] were used to quantify nestedness of biadjacency matrices. NODF is based on two characteristics of nestedness: decreasing fill and paired overlap. Because NODF depends on the order of rows and columns, we first sort the biadjacency matrix by decreasing row and column degrees to maximize the NODF score [8]. NODF is defined as

$$NODF = \frac{\sum N_{paired}}{[\frac{n(n-1)}{2}] + [\frac{m(m-1}{2}]}, \tag{S6}$$

where $n$ is the number of columns and $m$ is the number of rows in the matrix. Let row $i$ be located above row $j$, and column $k$ to the left of column $l$. For any pair of rows i and j, $N_{paired} = 0$ if the marginal total of row j is greater than or equal to that of row i. For any pair of columns $k$ and $l$, $N_{paired} = 0$ if the marginal total of column $l$ is greater than or equal to that of column $k$. Else, $N_{paired}$ is equivalent to the paired overlap ($PO$). For a pair of rows, $PO_{ij}$ is the percentage of 1s in row $j$ that are found in an identical column position in row $i$. For a pair of columns, $PO_{lk}$ is the percentage of 1s in column $k$ that are found in an identical row position in column $l$.

NODF scores are highly dependent on interaction density [7], so care was taken to avoid directly comparing scores between networks that differ substantially in their number of edges. In our simulations, the decline in probability of a nested interaction is matched exactly by an increase in the probability of a non-nested interaction, maintaining the same expected interaction density across the biadjacency matrix. This results in the observed homogeneity in null distributions for interaction network NODF scores (Figure 4). Co-occurrence networks, by contrast, do not have the same consistency in null NODF distributions, as they vary in edge density across the ranges of nestedness. Similar to the nested bipartite networks, interaction densities in the biadjacency matrix were kept constant while increasing decreasing the degree of modularity. The expected number of interactions per species was kept constant when varying the number of modules.

**Modularity:** Barber's Modularity $Q_b$, a bipartite version of the unipartite modularity introduced by Newman and Girvan [9, 10], was used to quantify modularity of the biadjacency matrices. Modularity $Q_b$ is formally defined as

$$Q_b = \frac{1}{2m} \sum_{i,j} (A_{ij} - P_{ij}) \delta(g_i, g_j), \tag{S7}$$

where $A_{ij}$ are the adjacency matrix elements, $P_{ij}$ is a probability in the null model that an edge exists between species $i$ and $j$, and $\delta$ is the Kronecker delta function. $\delta(g_i, g_j) = 1$ when species $i$ and $j$ are in the same module, and 0 otherwise. In practice, we translate the modularity metric as it is implemented in the MATLAB BiMat library into a Python function (see supplemental code) [11]:

$$Q_b = \frac{1}{E} \sum_{i,j} (B_{ij} - \frac{k_i d_j}{E}) \delta(g_i, h_j), \tag{S8}$$

where $B_{ij}$ are elements of the biadjacency matrix, $g_i$ and $h_i$ are the module indexes of nodes $i$ (that belong to class $V_1$) and $j$ (that belong to class $V_2$), $k_i$ is the degree of node $i$, $d_j$ is the degree of node $j$, and $E$ is the number of edges in the network.

Unlike nestedness, which can be calculated as a property of the biadjacency matrix as a whole, modularity by definition requires a concept of 'within' vs 'between' modules. Therefore, calculating $Q_b$ requires a given node partition. In each simulation, $Q_b$ reflects the modularity score of an optimal partition determined by the BRIM algorithm, implemented with the CDlib Python package [12]. By determining optimal node partitions for an underlying interaction network and its resulting co-occurrence network, we assessed clustering correctness similarly to the unipartite case. A pair of species within the same class was defined as correctly clustered if the two species were either clustered together (in the same module) in both the interaction and co-occurrence networks (a true positive), or *not* clustered together in both the interaction and co-occurrence networks (a true negative).

**Null models**: Both nestedness and modularity scores scale with other network attributes and are thus not necessarily comparable across networks. Therefore, NODF and $Q_b$ scores were calculated for 1000 random matrices with the same edge density as a given interaction or co-occurrence biadjacency matrix (obtained by randomly permuting the edges of the biadjacency matrix). The biadjacency matrix was considered significantly nested (modular) if its NODF ($Q_b$) score fell above (significantly anti-nested/modular, if below) the 95% confidence interval based on these 1000 randomizations.

For nested host-parasite networks, we found that statistically significant nestedness arose in co-occurrence networks even when the generative interaction networks were not significantly nested. Because nestedness is driven by overall fill [7], heterogeneity of degree distributions [13], and proper subsetting of specialist interactions within generalist interactions, we hypothesized that host-parasite interactions may produce unexpectedly heterogeneous degree sequences in co-occurrence networks, but not properly subset interactions. To test this hypothesis, we used an alternate, partially constrained null model (Probable Rows and Columns; PRC [14]) in which, for any matrix entry $a_{ij}$,

$$P(a_{ij} = 1) = (\frac{P_i}{n} + \frac{P_j}{m})/2, \tag{S9}$$

where $P_i$ is the marginal total of row $i$ of the given biadjacency matrix, $P_j$ is the marginal total of column $j$ in the given biadjacency matrix, $n$ is the number of columns, and $m$ is the number of rows. Else, $a_{ij} = 0$. In this way, the expected total number of edges and marginal totals remain consistent with the biadjacency matrix. Deviations from the PRC null more directly represent the subsetting of relatively more specialist interactions within relatively more generalist interactions.

##### S1.5.3 Degree distributions

Degree distributions were fitted and evaluated using the R package poweRlaw [15, 16]. This package implements comparisons between candidate distributions using Vuong's test, a likelihood ratio test for non-nested hypotheses which is a statistically robust way to identify relationships that are likely to be power laws [17, 18]. The candidate distributions that were evaluated were power law, log-normal, exponential, and Poisson. All distributions were fitted only for degree greater than the $xmin$ parameter, which is inferred when fitting the power law distribution, as recommended by [15].

The expected statistical distribution of node degrees in an Erdos-Renyi(n, p) random graph is approximately a binomial distribution with parameters $S-1$ (the maximum node degree) and $p$, where $p = 1-(1-p^{(rand)})^2$ for an undirected graph, which is the probability that there is an edge in either direction from one node to the other (Note: this converges to a Poisson distribution when $S$ is large and $p$ is small). Deviations from this expected distribution arise because the expected degree of one node is not independent of the degree of the other nodes. For example, if the degree of node 1 is $S-1$ (it is connected to every other node) then the degree of node 2 cannot be 0, and is therefore dependent on the degree of node 1. However, when a small proportion of the possible edges are realized, this deviation is negligible.

In order to test whether we can reject the random network degree distribution expectation (binomial), we performed Chi-squared goodness of fit tests on binned degree distributions. We set the binomial parameters ($n$ and $p$) to the number of species minus 1 (the maximum degree of any node) and the maximum likelihood estimate respectively. In other words, let $d_i$ be the observed degrees of the nodes in the network. Then $H_0$: $d_i \sim Binom(n, \hat{p})$, where $n = S-1$ and $\hat{p}$ is the maximum likelihood binomial probability given the observed degree distribution. We then computed the expected number of nodes in each degree bin under the maximum likelihood binomial distribution, and compare this to the observed numbers of nodes in each bin. The bins were computed by initially splitting the distribution into bins of length 1, and then collapsing the bins together until all bins have at least 5 expected counts. This means that the bin lengths will be longer in the tail of the distribution. The null hypothesis that the observed counts come from a binomial distribution is rejected when the p-value of the Chi-squared test is less than 0.05. The degrees of freedom of the Chi-squared test was set to the number of bins minus 2 (subtract an extra degree of freedom due to inferring the maximum likelihood parameter).

##### S1.6 Code availability

All code is available at `https://github.com/Fiona-MC/TheoryCoOccur-pub`.

#### S2 Other Supplemental Tables

##### S2.1 Power law minimum samples

Table S2.1 Caption: Over 10,000 samples drawn directly from a power law disribution are needed to confidently detect the difference between candidate distributions using Vuong's test. Here, samples were drawn directly from a discrete power law distribution as $X \sim PowerLaw(X_{min} = 2, \alpha = 3)$ using the R package PoweRlaw [15], from a Binomial distribution as $X \sim Binom(100, 0.2)$, from a Poisson distribution as $X \sim Poisson(20)$ and from a Log Normal distribution as $X \sim LogNormal(0, 3)$. Then PoweRlaw [15] was used to test the data for different $n \in \{50, 100, 200, 500, 1000, 10000, 100000, 1000000\}$ ten times for each $n$ value. All four distributions were compared for all data and a conclusion was made only if one distribution was a statistically significantly better (p¡0.05) fit than all of the others. Note that these are of course not all of the distributions that could be tested, so it is hard to conclude for certain that data comes from a particular distribution. Columns:

- `true_distribution`: the distribution the data was drawn from.
- `n`: the number of (iid) samples from the distribution.
- `n_replicates`: the number of times this experiment was repeated to get mean p- and R-values
- `mean_R_dist1_vs_dist2`: R statistic from Vuong's test comparing the maximum likelihood fit for distribution 1 and 2 (plaw = Power Law, pois = Poisson, lnorm = Log Normal, exp = Exponential).
- `mean_p_dist1_vs_dist2`: p-value from Vuong's test comparing the maximum likelihood fit for distribution 1 and 2 (plaw = Power Law, pois = Poisson, lnorm = Log Normal, exp = Exponential).
- `proportion_correct`: Proportion of the 10 replicates where we could confidently conclude that the correct distribution was a better fit than all of the others.

#### S2.2 PowRlaw

Table S2.2 caption: Sheet 1: Results for attempting to infer the degree distribution for 10 interaction and their corresponding co-occurrence networks for each degree distribution type: power law, exponential, and binomial. For each, we report the p-value for the goodness of fit test. If significant, this test indicates that the distribution is NOT a power law. Also reports the R statistics for Vuong's test comparing an exponential fit, a power law fit, a log normal fit, and a Poisson fit to the observed degree distributions. A positive R indicates that the Power Law fit is better than the other fit, and the p-value indicates statistical significance of that difference.

Sheet 2: Averages of the data in Sheet 1 across the 10 replicates.

Variables:

- `degDistType`: type of distribution that the original degree distribution of the interaction network follows.
- `p_gof_cooc` p-value for goodness of fit test for the null hypothesis that the distribution is a power law for the co-occurrence network degree distribution.
- `R_exp_cooc` R statistic for Vuong's test for a comparison of the exponential and power law distributions for the co-occurrence network.
- `p_exp_cooc` p-value for Vuong's test for a comparison of the exponential and power law distributions for the co-occurrence network.
- `R_logn_cooc` R statistic for Vuong's test for a comparison of the log normal and power law for the co-occurrence network.
- `p_logn_cooc` p-value for Vuong's test for a comparison of the log normal and power law distributions for the co-occurrence network.
- `R_pois_cooc` R statistic for Vuong's test for a comparison of the Poisson and power law for the co-occurrence network.
- `p_pois_cooc` p-value for Vuong's test for a comparison of the Poisson and power law distributions for the co-occurrence network.
- `p_gof_inter` p-value for goodness of fit test for the null hypothesis that the distribution is a power law for the interaction network degree distribution.

- `R_exp_inter` R statistic for Vuong's test for a comparison of the exponential and power law distributions for the interaction network.
- `p_exp_inter` p-value for Vuong's test for a comparison of the exponential and power law distributions for the interaction network.
- `R_logn_inter` R statistic for Vuong's test for a comparison of the log normal and power law for the interaction network.
- `p_logn_inter` p-value for Vuong's test for a comparison of the log normal and power law distributions for the interaction network.
- `R_pois_inter` R statistic for Vuong's test for a comparison of the Poisson and power law for the interaction network.
- `p_pois_inter` p-value for Vuong's test for a comparison of the Poisson and power law distributions for the interaction network.
- `cooc_alpha` Maximum likelihood $\alpha$ parameter for the co-occurrence network degree distribution power law fit.
- `cooc_xmin` Maximum likelihood $x_{min}$ parameter for the co-occurrence network degree distribution power law fit.
- `inter_alpha` Maximum likelihood $\alpha$ parameter for the interaction network degree distribution power law fit.
- `inter_xmin` Maximum likelihood $x_{min}$ parameter for the interaction network degree distribution power law fit.
- `correct_cooc` 1 if the correct distribution was inferred for the co-occurrence network.
- `correct_inter` 1 if the correct distribution was inferred for the interaction network.

#### S2.3 GoF test for binomial

Caption for table S2.3: Results of a goodness of fit of the degree distributions for interaction and co-occurrence networks. Null hypothesis for all tests is that the data is drawn from $x \sim Binomial(N, p)$

where N is the number of nodes and p is the maximum likelihood parameter for the binomial distribution given the observed node degrees. Sheet 1 shows the summary of results for the co-occurrence networks. Sheet 2 shows the full results for the same co-occurrence network tests. Sheet 3 shows the summary of results for the interaction networks. Sheet 4 shows the full results for the same interaction network tests. Tests were run for 50 replicates for interaction strength $F_{max} \in \{0.1, 0.2, 0.33\}$, number of species $S = 1000$, and probability of positive interaction $q = 0.001$ for exponential, binomial, and power law starting interaction network degree distributions. Additionally they were run for 100 replicates for interaction strength $F_{max} \in \{0.1, 0.2, 0.33\}$, number of species $S = 100$, and probability of positive interaction $q = 0.5$ for exponential ("exp"), binomial ("rand"), and power law ("power") starting interaction network degree distributions.

### S3 Frequency of interaction types

For ecological interactions, each type of interaction can be understood at a combination of the signs of the interaction coefficients between the two species. For example, a mutualism would be an interaction with positive coefficients in both directions, whereas competition would have negative coefficients in both directions, and trophic or parasitic interactions would have a positive coefficient in one direction and a negative coefficient in the other direction. However, for some of these, it is unclear whether the interaction would induce positive or negative correlation [19], and in any case since correlation is symmetric, it is impossible to have an edge in the co-occurrence matrix that is positive in one direction and negative in the other.

Although we cannot directly compare the types of edges in the network, we can compare the correspondence between the proportion of positive and negative entries in the interaction versus co-occurrence matrices. We find that without environmental confounding, the proportion of positive edges in the interaction network is strongly correlated with the proportion of significant positive correlations in the co-occurrence network (Figure S1). However, while the interaction network proportion of positive edges ranges from 0 to 0.8, the co-occurrence proportion only ranges from 0.4-0.55. At lower interaction strengths, the trend is less clear.

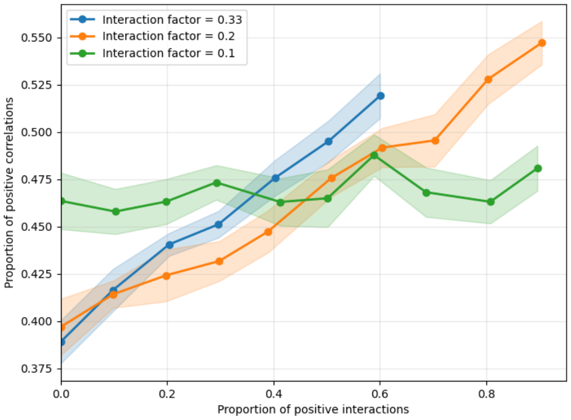


Figure S1: Proportion of positive edges (number of positive edges / total number of edges) in co-occurrence network versus proportion of positive edges in interaction network (number of positive interactions / total number of interactions) for $F_{max} = 0.33$, 0.2, and 0.1

### S4 Varying species' carrying capacities

To verify that results were robust to variation in species' carrying capacities, self-limitation parameter $\alpha$ was varied. For $\alpha \sim Uniform(0.00067, 0.002)$, corresponding to carrying capacities uniformly distributed between 500 and 1500, there was no qualitative change in the relationship between interaction strength and resulting Pearson correlations (Fig. S2A), the relationship between the proportion of positive interactions and the proportion of positive edges in the co-occurrence network (Fig. S2B), or the degree to which modular structures in an interaction network are retained in the co-occurrence network (Fig. S2C).

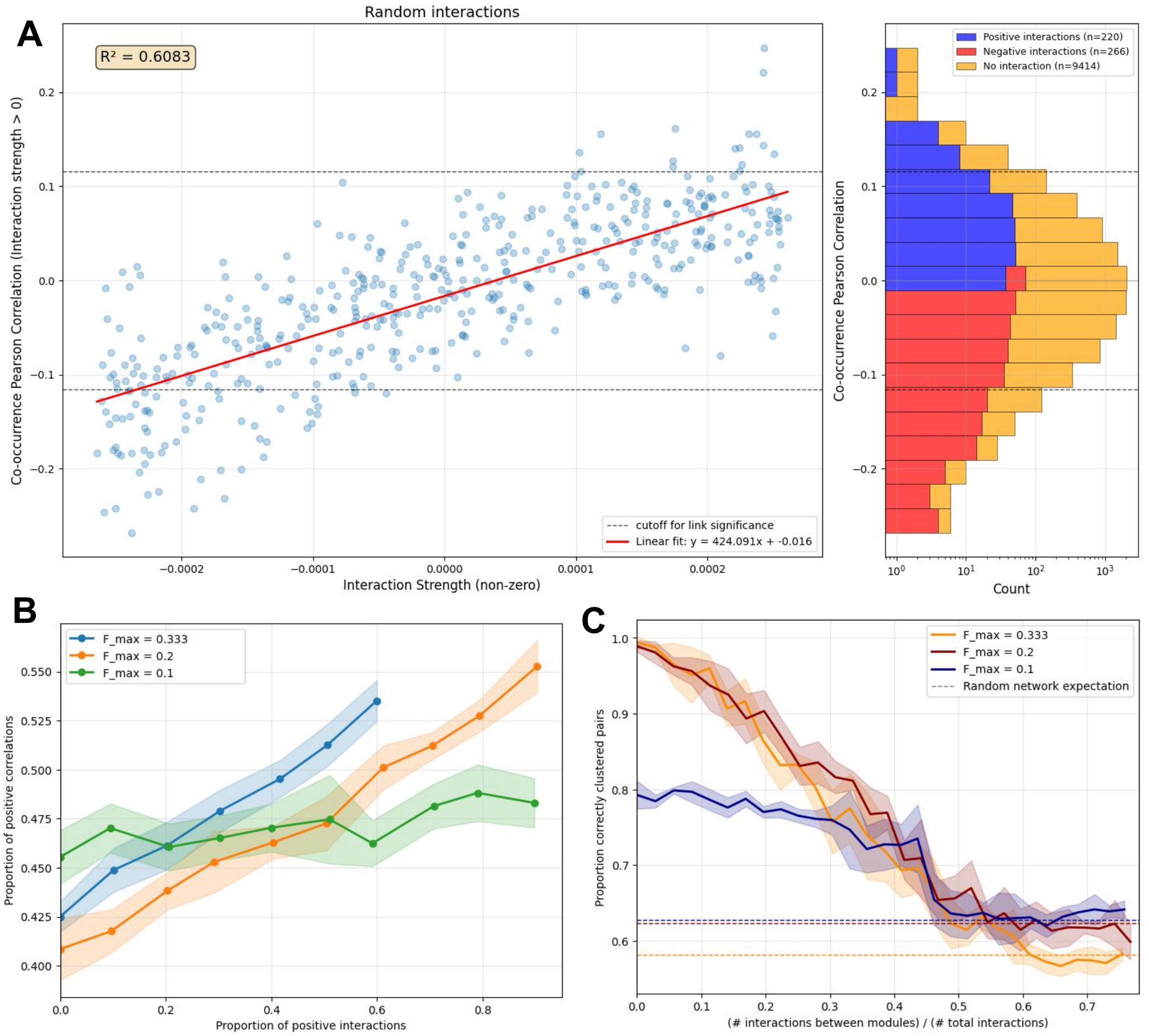


Figure S2: **Variable carrying capacities.** A, B, and C are the counterparts of Figure 1 in the main text, Figure S1, and Figure 2D in the main text, respectively, with $\alpha \in [0.00067, 0.002]$ rather than $\alpha = 0.001$. A) Co-occurrence network Pearson correlation versus interaction strength and distribution of co-occurrence network Pearson correlations for interacting and non-interacting species (log scale) from an interaction network formed by an Erdos-Renyi random graph. B) Proportion of positive edges in co-occurrence network versus proportion of positive edges in interaction network. C) Proportion of pairs of nodes that are correctly clustered versus the proportion of the realized edges in the interaction network that are between modules. A pair of nodes is defined as "correctly clustered" if both nodes are in the same cluster in the interaction network and in the co-occurrence network, or if the two nodes are in different clusters in both the interaction and co-occurrence networks. Error bars are shown for the standard deviation of five replicates. Random network expectations are shown for the same metric computed for an Erdos-Renyi random graph with the same number of edges and the same maximum interaction strengths.

### S5 Bipartite interaction networks

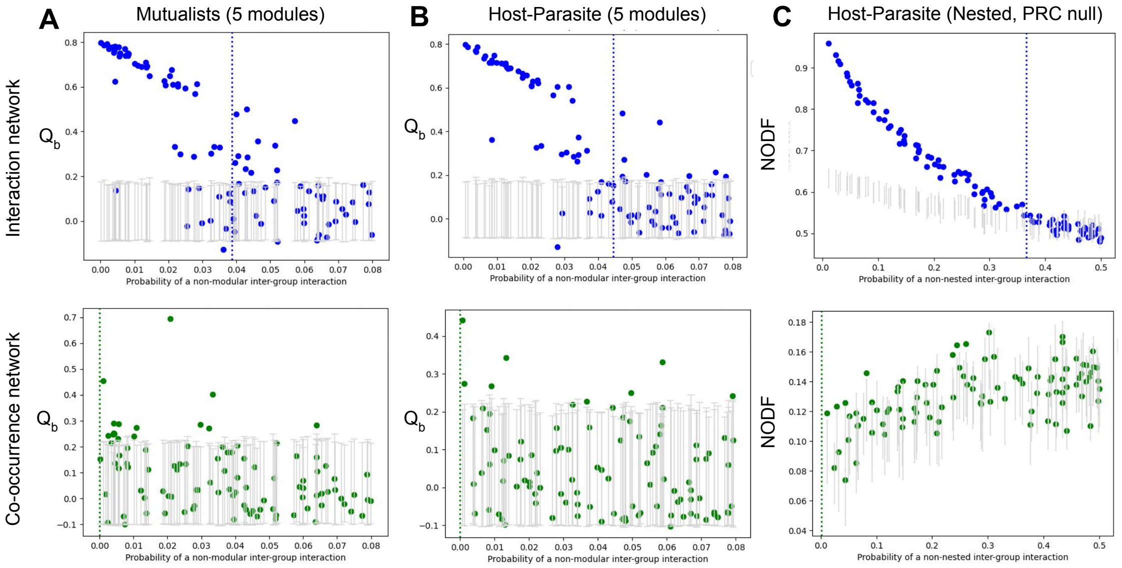


Figure S3: **Alternative structures and null models for bipartite interaction networks.** Interaction and co-occurrence networks of modular bipartite mutualist (A), host-parasite (B), and nested host-parasite (C) networks. $Q_b$ or NODF scores were calculated for the biadjacency matrices of both the interaction network and corresponding co-occurrence network. Vertical dotted lines indicate the $p = 0.5$ decision threshold for a binary encoding of the presence of statistically significant nestedness. In (A) and (B), grey bars represent 95% confidence intervals for $Q_b$ based on $10^3$ randomizations of the interaction or co-occurrence network. In C, grey bars represent 95% confidence intervals for $Q_b$ based on $10^3$ partially constrained randomizations (PRC null model) of the interaction or co-occurrence network. **(A)** Inter-class mutualist interactions are structured with varying degrees of modularity as described in S1.2, with $p^{(m)} \leq 0.5$ to ensure the per-species edge density remains consistent with the 10 module case. All inter-class interaction coefficients $a_{ij} > 0$. **(B)** Inter-class host-parasite interactions are structured with varying degrees of modularity as in (A). For a given host-parasite interaction between host $i$ and parasite $j$, $a_{ij} < 0, a_{ji} > 0$. **(C)** Inter-class host-parasite interactions are structured with varying degrees of nestedness as described in S1.2.

### S6 Indirect interactions can reinforce modules

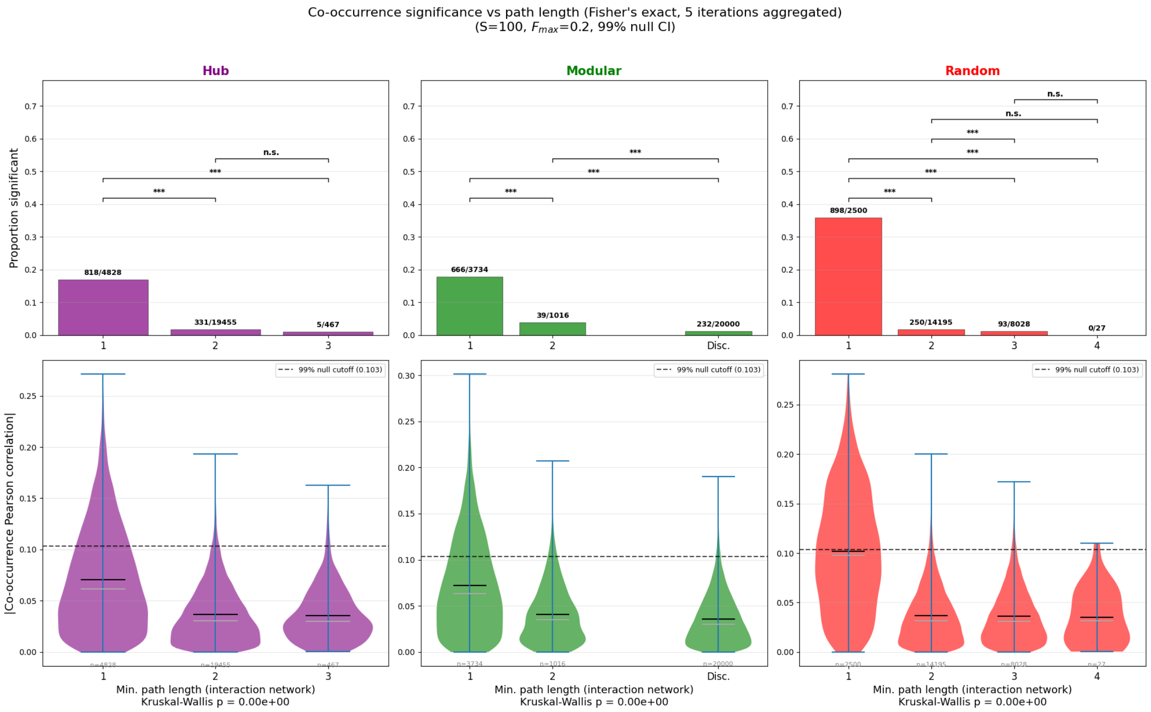


Figure S4: Top row: Proportion of significant co-occurrence edges (Number of edges / Number of possible edges) that are distance 1, 2, or 3 apart in the interaction network (ie distance 1 is a direct interaction edge). Bottom row: Distribution of the absolute value of co-occurrence Pearson correlations for pairs of taxa that are distance 1, 2, or 3 apart in the interaction network.

### S7 Role of the environment

Many of the patterns seen in co-occurrence networks that are caused by interaction patterns can also be caused by shared responses to the environment. For example, modules may be formed by positive associations when species respond the same way to environments. Co-occurrence modules formed in this way will be much more pronounced if only positive co-occurrence edges are considered, and many negative edges will exist between modules. For species with uniform trait values with two available environments, we find that species with more similar trait values have significantly higher co-occurrence correlations. However, when we form modules, they do not fall along the lines of environmental associations because there are many negative associations between modules. If we filter associations for positive correlations, there is a clear signal indicating that there is an enrichment of positive edges within modules and negative edges between modules (Supplementary figure S5). Additionally, we see that when modules are formed by inter-specific interactions, the modules are enriched for both positive and negative interactions. On the other hand, when the modules are formed by environmental preference, the modules are enriched for positive co-occurrence edges, while there are more negative edges between the modules compared to within. This may suggest ways to predict whether modules are likely primarily driven by the environment or by inter-species interactions in empirical co-occurrence data.

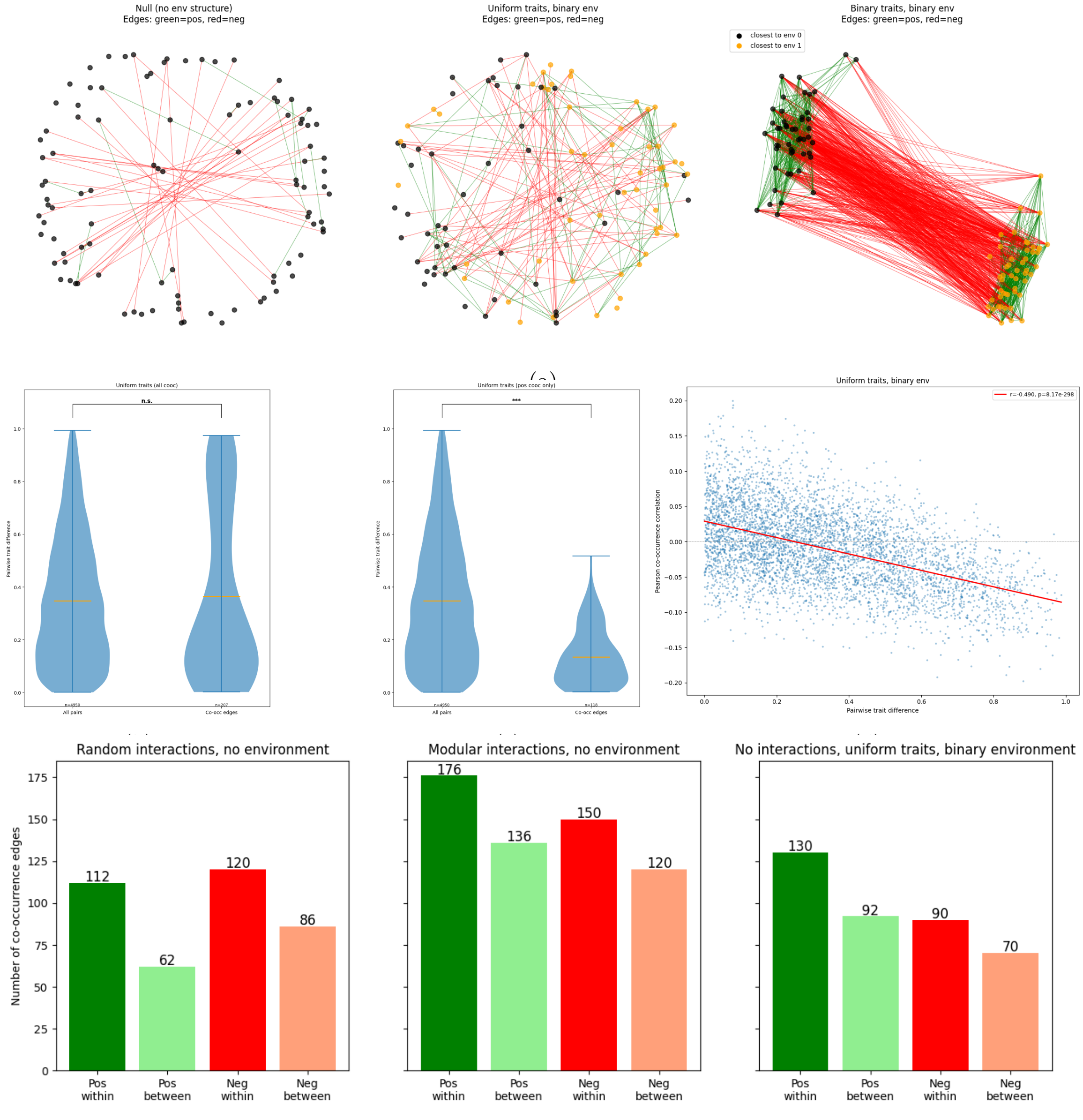


(e)
Figure S5

### S8 Comment on keystoneness

Species with disproportionately large numbers of interactions are sometimes interpreted as "keystone species", but we avoid that term here because it has been defined in multiple ways [20]. Different forms or definitions of keystoneness may be correlated with different properties of the network. For example, we find that degree centrality and betweenness centrality in the co-occurrence network predict high-degree nodes in the interaction network, which we call hubs. In a similar simulation framework, when keystoneness was defined as the effect of species removal on species richness, high co-occurrence network node degree, high co-occurrence closeness centrality, low co-occurrence betweenness centrality, and high co-occurrence transitivity are good predictors of high keystoneness, while these properties in the interaction network have no predictive power [1]. This is a direct consequence of the definition of keystone species because we define hubs based on the degree centrality in the interaction network, whereas Berry and Widder define keystoneness based on the results of the simulation, which has a close relationship with co-occurrence correlation. Different metrics of centrality in both networks may help to understand different aspects of keystoneness, and therefore it is essential to carefully define the keystone concept in each case.

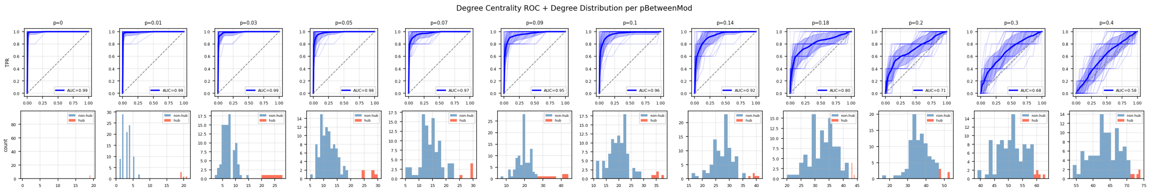


Figure S6: ROC curves for degree centrality and degree distributions for hub networks starting at different levels of background interactions ($p_{ij} = 0$ to $p_{ij} = 0.4$).

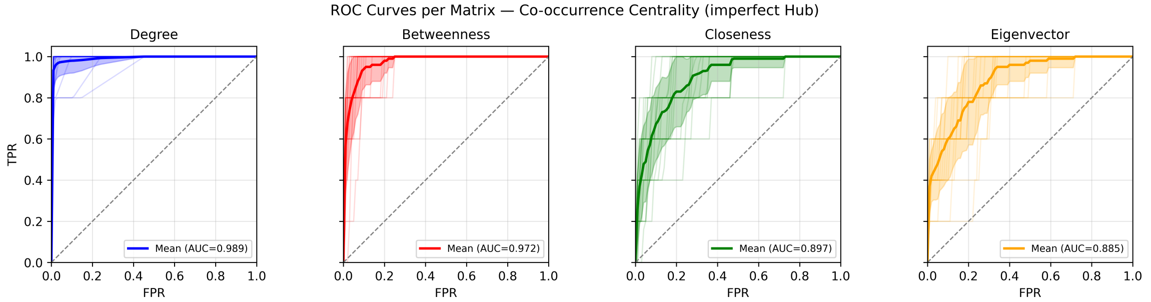


Figure S7: (a) ROC curves for predicting hub nodes from co-occurrence data using various centrality metrics when the maximum interaction strength is $F_{max} = 1/5$. Plot shows average results of 20 simulations with 100 species each, where each simulation includes 5 hub nodes. Shaded error bars are one standard deviation across the replicates, and single runs are depicted as lighter lines. 90% of the interactions between the hub and other species in the module are realized. 1% of the interactions between other pairs are realized.

### S9 Skewed Null Distribution of Correlations

The way our simulation is constructed causes the correlation values to be slightly negatively skewed, even without interactions. This is because we sample the species without replacement, leading to their presences being non-independent even in a non-interaction case. The way we construct the null distribution (running the simulation without interactions) accounts for this. Using a permutation-based null does not account for this and therefore was not used (Supplementary Figure S8).

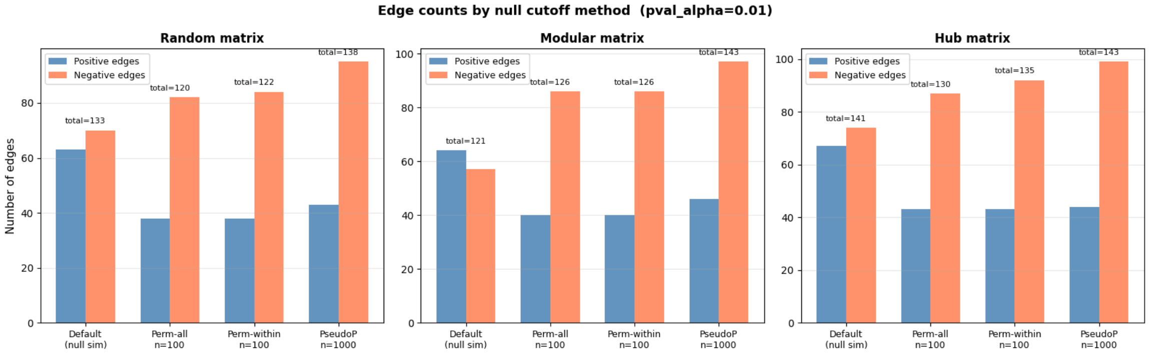


Figure S8: Numbers of significant positive and negative edges in the co-occurrence matrix using different methods of constructing a null. The Default is the method described in this paper, and the others are all permutation-based tests.

Specifically:

In general, for events A and B,

$\mathbb{P}(A, B) = \mathbb{P}(A)\mathbb{P}(B|A)$

If we let event A be presence of one species and B be presence of another (and $\bar{A}$ and $\bar{B}$ be absence of species A and B respectively), then under the null hypothesis that there is no interaction:

$\mathbb{P}(A) = \mathbb{P}(B) = 50/100$

$\mathbb{P}(\bar{A}) = \mathbb{P}(\bar{B}) = 50/100$

$\mathbb{P}(B|A) = 49/99$

$\mathbb{P}(B|\bar{A}) = 50/99$

$\mathbb{P}(A|\bar{B}) = 50/99$

$\mathbb{P}(\bar{B}|\bar{A}) = 49/99$

So then:

$\mathbb{P}(A, B) = 0.247$

$\mathbb{P}(\bar{A}, \bar{B}) = 0.247$

$\mathbb{P}(\bar{A}, B) = 0.253$

$\mathbb{P}(A, \bar{B}) = 0.253$

In other words, sampling species without replacement makes it slightly more likely that we get negative correlations as opposed to positive even when there is no interaction.

### S10 Additional parameter combinations

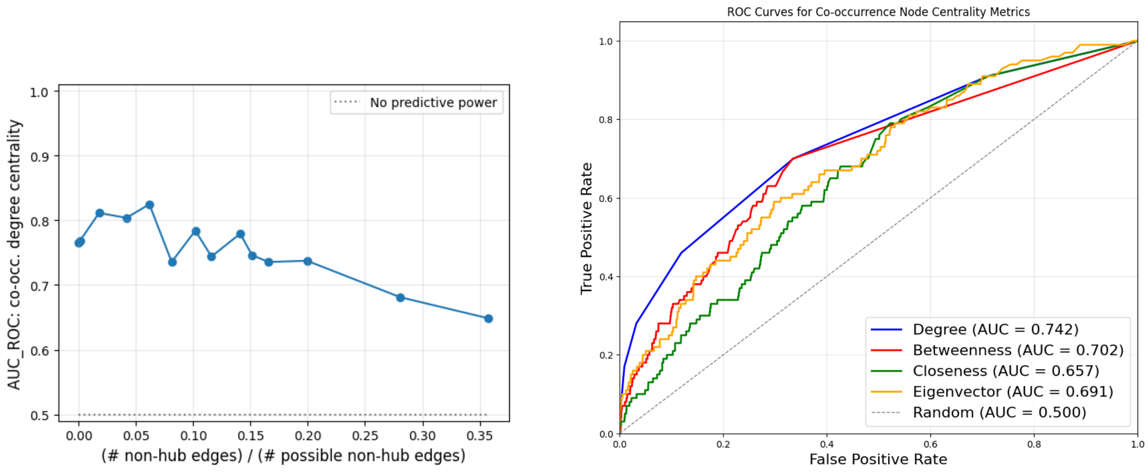


(a) (b)

Figure S9: Same as main text hub figures but for $F_{max} = 1/10$

To illustrate one reason why we get such a high false discovery rate for direct interactions in this setting where a relatively low proportion of possible edges are realized (i.e. interactions are sparse), imagine trying to identify just one interaction among 100 species. There are (100 choose 2) = 4950 possible pairs, so if you use a 95% confidence set, you would expect that you would find over 200 significant correlations that don't represent direct interactions. Even if you are more conservative than that, it is likely that by chance some correlations between non-interacting species will be as strong or stronger than the interacting pair.

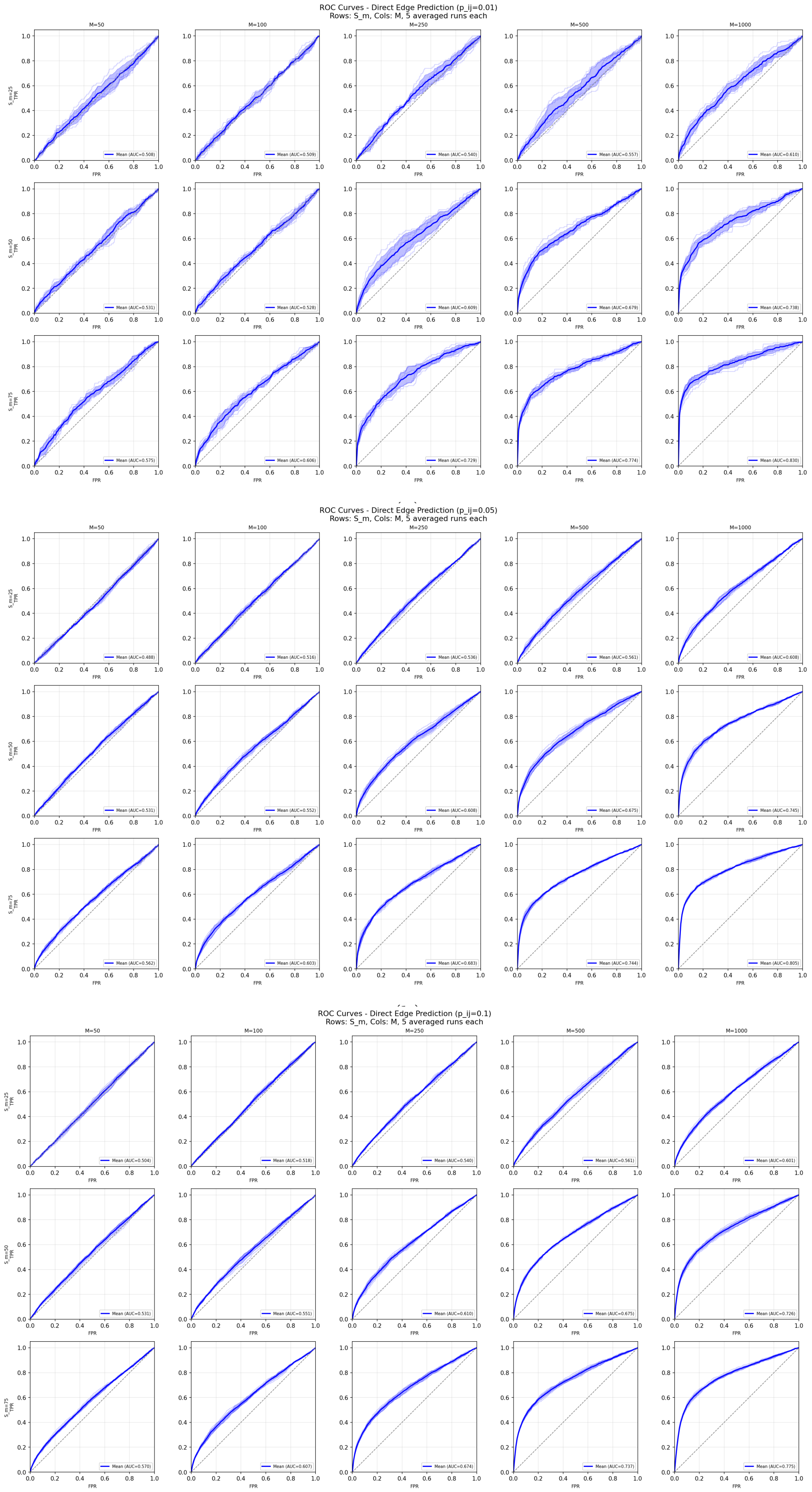


(c)

Figure S10: Direct edge prediction with different parameters.
